# Pangenome evolution in environmentally transmitted symbionts of deep-sea mussels is governed by vertical inheritance

**DOI:** 10.1101/2021.11.12.468352

**Authors:** Devani Romero Picazo, Almut Werner, Tal Dagan, Anne Kupczok

## Abstract

Microbial pangenomes vary across species; their size and structure are determined by genetic diversity within the population and by gene loss and horizontal gene transfer (HGT). Many bacteria are associated with eukaryotic hosts where the host colonization dynamics may impact bacterial genome evolution. Host-associated lifestyle has been recognized as a barrier to HGT in parentally transmitted bacteria. However, pangenome evolution of environmentally acquired symbionts remains understudied, often due to limitations in symbiont cultivation. Using high-resolution metagenomics, here we study pangenome evolution of two co-occurring endosymbionts inhabiting *Bathymodiolus brooksi* mussels from a single cold seep. The symbionts, sulfur-oxidizing (SOX) and methane-oxidizing (MOX) gamma-proteobacteria, are environmentally acquired at an early developmental stage and individual mussels may harbor multiple strains of each species. We found differences in the accessory gene content of both symbionts across individual mussels, which are reflected by differences in symbiont strain composition. Compared to core genes, accessory genes are enriched in functions involved in genome integrity. We found no evidence for recent horizontal gene transfer between both symbionts. A comparison between the symbiont pangenomes revealed that the MOX population is less diverged and contains fewer accessory genes, supporting that the MOX association with *B. brooksi* is more recent than that of SOX. Our results show that the pangenomes of both symbionts evolved mainly by vertical inheritance. We conclude that environmentally transmitted symbionts that associate with individual hosts over their lifetime show features of narrow symbioses, where the frequency of HGT is constrained.

## Introduction

Bacterial populations can show enormous genomic diversity, which comprises nucleotide differences between homologous sequences and variation in the accessory gene content. In particular, gene content diversity is described by the species pangenome, which consists of all the genomic sequences present across individuals of a bacterial species. The core genes in a pangenome are present in each individual while the remaining genes are considered accessory (Brockhurst et al. 2019). Pangenome size and structure vary across bacterial species (Maistrenko et al. 2020) and pangenomic diversity is important for bacterial adaptation in environmental species, where accessory genes are often niche-specific, e.g., for the marine *Prochlorococcus* (Kashtan et al. 2014). To understand microbial adaptation, it is thus crucial to understand the evolutionary processes that shape pangenome diversity. The main processes that give rise to microbial pangenomes are gene duplication and loss during vertical inheritance as well as gene acquisition via horizontal gene transfer (HGT). HGT enables the transfer of genetic material between microbial individuals that are not related by inheritance (Hall et al. 2017) and is particularly relevant for the evolution of microbial pangenomes (Treangen & Rocha 2011; Tria & Martin 2021). Some mechanisms of HGT involve the activity of mobile genetic elements (MGEs) - such as phages, plasmids, or transposons - for transferring genetic material between different DNA strands.

Many bacterial species are known to have a host-associated lifestyle, i.e., they are strictly or facultatively associated with eukaryotic hosts. Symbionts have different modes of transmission; parentally (or vertically) transmitted bacteria are transferred from adults to their progeny, while environmentally (or horizontally) transmitted bacteria are acquired from the environment either from a free-living population or other hosts. Mixed transmission modes are also common over long evolutionary time (Bright & Bulgheresi 2010; Russell 2019). The host association has important implications for the adaptation of symbionts via HGT since bacterial populations that share a habitat may be able to access the habitat-specific gene pool by HGT (Bordenstein & Reznikoff 2005; Newton & Bordenstein 2011; Polz et al. 2013). Indeed, several studies demonstrated that gene transfer from locally adapted populations may facilitate host colonization. For example, in plant-associated communities, MGEs enabled the adaptation of locally adapted nitrogen-fixing soil bacteria to associate with novel crops during their domestication (Greenlon et al. 2019) and in sponges, diverse functions that potentially provide a selective advantage to the symbionts in that niche were acquired by HGT (Robbins et al. 2021). Notably, these examples stem from environmentally transmitted symbionts that might have a wider potential for HGT compared to parentally transmitted symbionts. First, infection of a host by multiple symbionts results in a shared environment, where the chances for HGT are higher and second, genes can potentially be acquired from environmental bacteria during the free-living stage. Indeed, very few HGT events have been reported in well-studied insect symbioses, potentially due to genetic isolation linked to the intracellular lifestyle and parental transmission (Pinto-Carbó et al. 2016; López-Madrigal & Gil 2017; Waterworth et al. 2020).

Previous studies showed that the evolution of endosymbiont genomes is characterized by rare HGT and fewer accessory genes compared to environmental bacteria (Kloesges et al. 2011; Brockhurst et al. 2019). This conclusion was drawn based on a few model symbionts that have been cultivated and sequenced and that are mostly parentally transmitted. However, less is known about symbionts that are metabolically dependent on their host and hence cannot be well cultivated under laboratory conditions (Pande & Kost 2017) and were thus excluded from previous studies. In the last 20 years, cultivation-independent approaches, such as metagenomics, revolutionized our view of microbial diversity (e.g., (Giovannoni et al. 2014; Castelle & Banfield 2018)). Additionally, deeply sequenced metagenomes provide adequate datasets for studying variation within microbial species (Denef 2019; Rossum et al. 2020). Metagenomics approaches enable us to assess the variation of all organisms in a particular environment. This approach revealed abundant strain diversity in symbionts, e.g., in the gut microbiome of humans and bees (Ellegaard & Engel 2019; Garud et al. 2019).

The presence of strain diversity has recently been reported for environmentally transmitted symbionts that reside in *Bathymodiolus* mussels. These symbionts are acquired during the mussels’ metamorphosis from a planktonic to a benthic lifestyle and are hosted in bacteriocytes within the gill epithelium, where they provide the mussel with nutrition (Won et al. 2003; Franke et al. 2021). *Bathymodiolus* can be infected by two chemosynthetic symbiont species, sulfur-oxidizing (SOX) and methane-oxidizing (MOX) gamma-proteobacteria. Although most *Bathymodiolus* species harbor only a single 16S phylotype for each symbiont, metagenomic analyses of multiple *Bathymodiolus* species showed that different SOX and MOX strains can be present within an individual mussel (Ansorge et al. 2019; Romero Picazo et al. 2019). An important role of MGEs and HGT in the evolution of SOX symbiont genomes from hydrothermal vents at the mid-atlantic ridge has been suggested. There, SOX genomes were found to contain high numbers of transposases, integrases, restriction-modification systems, and toxin-related genes, where the latter are also linked to MGEs (Sayavedra et al. 2015). In addition, it has been observed that co-occurring SOX strains from these sites differ in the content of genes involved in energy and nutrient utilization and viral defense mechanisms (Ansorge et al. 2019).

To study the evolution of pangenomes in symbiont populations, we analyzed the pangenomes of environmentally transmitted symbionts that reside in closely related, nearby hosts. To this end, 19 *Bathymodiolus brooksi* mussels were sampled from a single location at a cold seep site in the northern Gulf of Mexico. Since *Bathymodiolus* symbionts cannot be cultured, high-resolution metagenomics data was collected by deeply sequencing homogenized gill tissue of each mussel (Romero Picazo et al. 2019). We previously obtained single-sample assemblies and used a gene-based binning approach to reconstruct the core genomes of SOX and MOX (Fig. 1). Based on single nucleotide variants (SNVs) within the core genes, reconstruction of core-genome-wide strains revealed eleven SOX strains that group into four clades, and six MOX strains that group into two clades (Romero Picazo et al. 2019). Mussel individuals may harbor one or multiple strains of each species. In particular, they can contain strains from one to three different SOX clades and from one or both MOX clades (Fig. 1 in (Romero Picazo et al. 2019)). We found a high variability of the nucleotide diversity π between samples, where samples with low π tend to have a lower strain diversity as estimated using α-diversity. We furthermore investigated genetic isolation using the fixation index F_ST_, which revealed generally high genetic isolation between samples and also clusters of low genetic isolation where samples show highly similar SNV states and frequencies, i.e., they contain similar populations for a particular symbiont. These clusters were also detected using the ecological measure β-diversity, thus they are also similar in strain composition and frequencies. Taken together, we found that the evolution of symbiont populations in individual mussels is characterized by genetic isolation, suggesting that symbionts are taken up at an early stage in the mussel life cycle and are then geographically isolated (Romero Picazo et al. 2019). Here, we study the effect of the geographical isolation on SOX and MOX pangenome evolution. To this end, we analyzed the population pangenomes of the SOX and MOX strains residing in these 19 mussels sampled from a single location.

**Figure 1.**
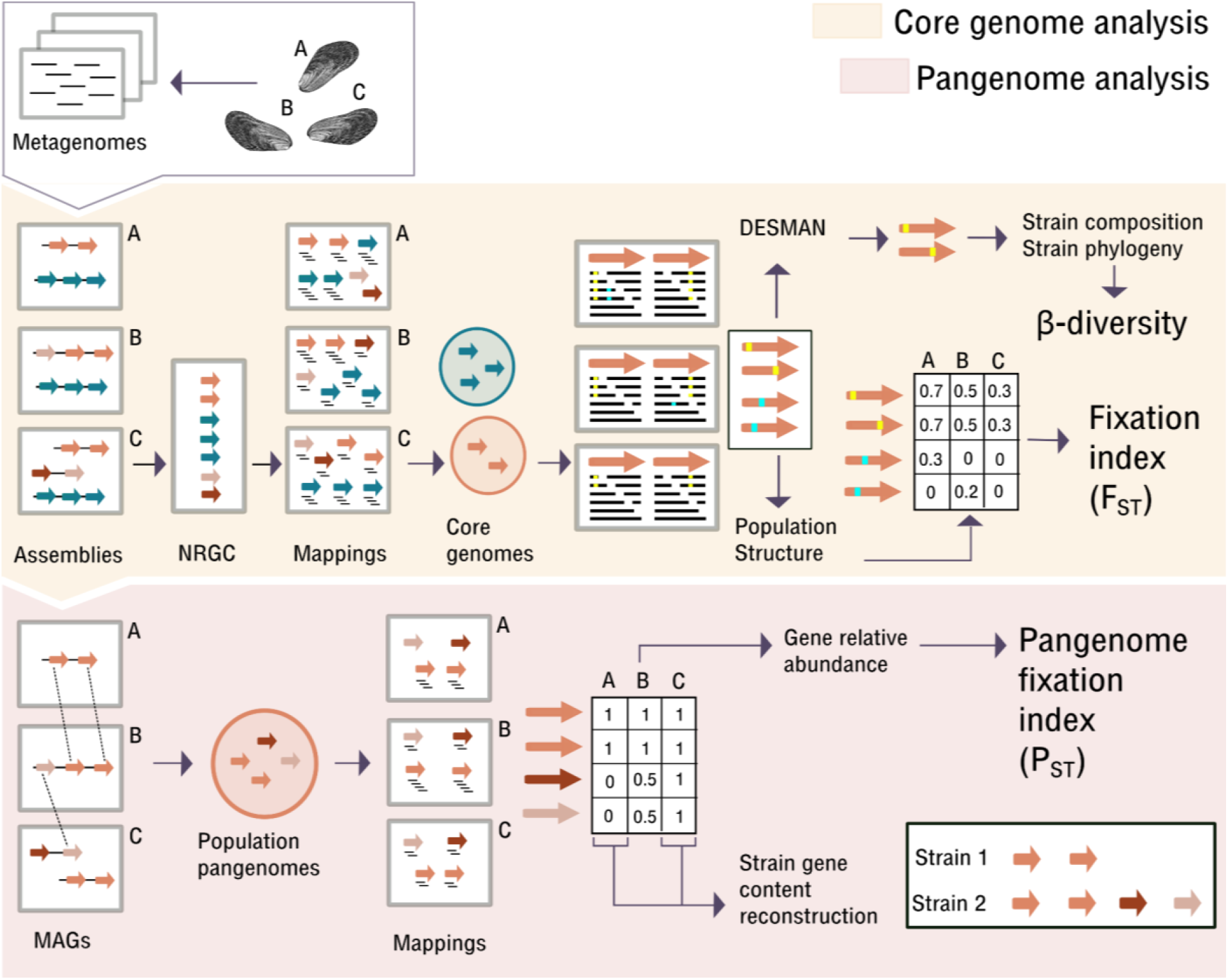
High-resolution metagenomic analysis workflow. Arrows represent ORFs (Open Reading Frames) inferred from the assemblies. Orange part: Core genome analysis as described before (Romero Picazo et al. 2019). The core genome analysis included the reconstruction of core genome strain sequences and estimation of population structure measured as both, β-diversity and F_ST_. Note that all SNVs are included in the F_ST_ calculation, whereas β-diversity is based on the strain composition and not all SNVs can be linked to strains by DESMAN. NRGC: non-redundant gene catalog. This catalog comprises gene cluster representatives obtained by grouping highly similar genes across samples. Red part: Pangenome analysis presented in this paper. The analysis is shown for a single species for simplicity. The network approach allows reconstructing population pangenomes. For each sample, the complete set of contigs containing genes from the species pangenome corresponds to the reconstructed MAG (metagenome-assembled genome). Using mapping, we estimated the coverages for all genes in the pangenomes in each sample, also including genes that have not been reconstructed on the contigs of that sample. The relative abundance of genes in the pangenome is then used to estimate P_ST_. Additionally, we reconstructed the gene content of single strains that are dominant in a sample.

## Results

### A network approach for the recovery of symbiont pangenomes from microbiome metagenomes

To reconstruct the SOX and MOX pangenomes, we recovered the accessory genomes using a network approach. The core genomes of both symbionts were used as starting points for the expansion of the pangenomes in the network. Here, we identified connections between the symbiont genes, which are located on different metagenomic contigs in each sample (Fig. 1). These connected contigs are therefore identified as related to one symbiont and correspond to the reconstructed metagenome-assembled genomes (MAGs). Our pangenome inference revealed clear differences between the two symbionts. The pangenome of the SOX population comprises 2,484 genes with a total length of 2.27Mbp. Of these, 962 (38.7% of total genes) were identified as accessory, where the majority are single-copy accessory genes (Table 1). Each SOX MAG contains between 1,640 and 2,055 genes (average 1,885) with genome lengths that range between 1.72 and 2.21 Mbp (average 2.06 Mbp) (Table S2A). The MOX population pangenome comprises 2,866 genes with a total length of 2.24 Mbp, where 414 (14.5% of total genes) are accessory and the majority of accessory genes are single-copy (Table 1). Each MOX MAG contains between 2,480 and 2,603 genes (average 2,546) with genome lengths between 2.39 and 2.50 Mbp (average 2.45 Mbp) (Table S2B). The SOX and MOX core genomes have an average GC content of 38% where the core GC content distributions differ significantly between SOX and MOX (Table 1, Fig. S1). In both symbionts, the GC content of the accessory genome is significantly lower than that of the core genome (Table 1, Fig. S1).

**Table 1:**
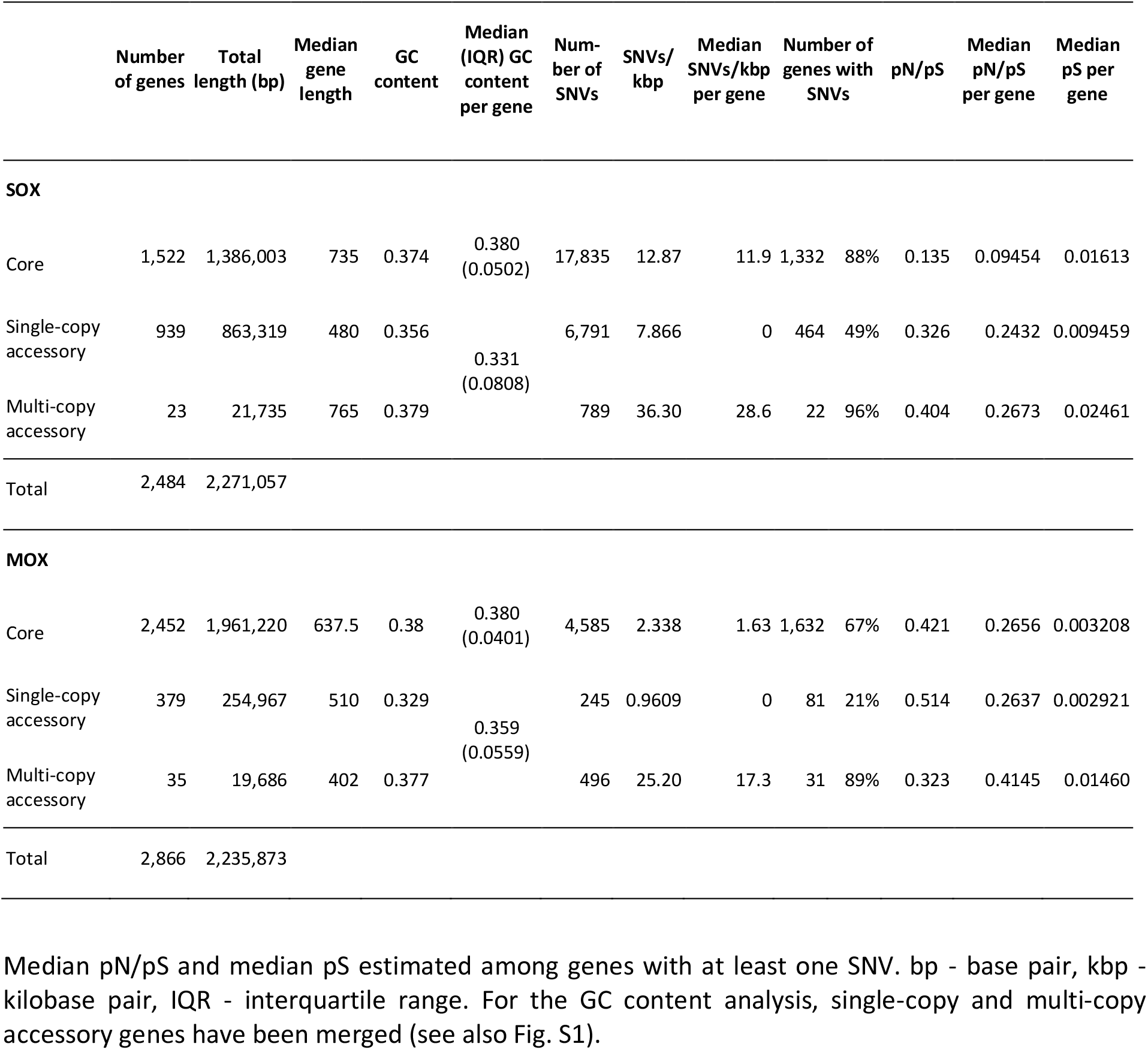
Description of SOX and MOX pangenomes.

To infer the reliability of our approach to reconstruct pangenomes from metagenomes, we investigated assembly statistics depending on the number of strain clades in each sample (Table S2, Fig. S3). We observed that a higher strain diversity results in longer MAGs and more genes for MOX supporting that the MAGs are population genomes including genes from multiple strains. In contrast, the length and number of genes on SOX MAGs does not increase with strain diversity, which indicates that contig fragmentation leads to missing regions in the MAGs. When mapping the samples on the SOX accessory genome, we detected more genes for samples with higher strain diversity; thus, samples with more strain clades indeed contain more accessory genes. Since the following analyses are based on the reconstructed pangenomes and the frequency of each gene in each sample, our approach provides a less biased assessment of gene presence in each sample compared to using MAGs.

Furthermore, to evaluate the sensitivity of the network approach to identify accessory genes, we estimated the recovery rate as the number of new genes that are added to the pangenome for each newly sampled mussel (Fig. S2A,B). We observed that 95% of the total accessory genome is detected when adding five samples for both bacterial species. This shows that our approach has the required sensitivity in order to recover accessory genes in the population pangenomes at this site. Sequencing coverage can further impact the sensitivity of our approach to recover pangenomes from metagenomic data. The two symbionts occur at different abundances within the mussel, which results in differences in the sequencing coverage of SOX and MOX. To investigate whether sequencing coverage might impact the sensitivity to detect accessory genes, we estimated the recovery rate for SOX when downsampling it to the MOX coverage. We found no decrease in the number of detected genes in SOX with this normalization, where also 95% of the total accessory genome is detected with five samples (Fig. S2C) and therefore the recovery of the accessory genome is comparable to the approach using the full coverage.

By comparing the pangenome characteristics of the two symbionts, we found that the MOX population pangenome contains more genes than SOX, whereas the accessory genome is larger in SOX. We found that the accessory genome sizes are less than 40% of the total pangenome sizes. Notably, this proportion is smaller than for the majority of known species pan-genomes, where 38 of the 43 species pangenomes estimated from RefSeq records (Ding et al. 2018) have an accessory genome fraction above 40% and the five exceptional species are all obligate pathogens (*Bordetella pertussis, Brucella mellitensis, Chlamydia trachomatis, Mycobacterium tuberculosis, Mycoplasma pneumoniae*). The low proportion of accessory genes in the pangenomes reconstructed here is likely due to the fact that we reconstruct the pangenomes from the sampling site, i.e., not that of the entire species, where the species pangenome is expected to be larger. The lower GC content in the accessory genomes of both species may indicate 1) that genes are transferred from an external source with low GC content, or that genes with lower GC content are preferably transferred or 2) that accessory and core genes are under different selection regimes, where the higher GC content in the core genome is maintained by purifying selection (Bohlin et al. 2017).

### Gene content shows genetic isolation between mussels that is explained by strain composition

The intracellular lifestyle of the symbionts results in strong geographic isolation between mussels and this leads to genetically isolated symbiont populations (Romero Picazo et al. 2019). To study how genetic isolation impacts pangenome evolution in the mussel symbionts, we examined the SOX and MOX gene content variation within and across individual mussels.

To analyze symbiont gene content diversity within individual mussels, we estimated the gene content diversity φ that is based on the relative frequency of genes in a population or subpopulation (see Methods). We found that φ is positively associated with the nucleotide diversity π and both measures increase with α-diversity estimated from the strain composition (Fig. 2A,B, S4A,B). We did not observe any difference in φ between SOX and MOX (two-sided Wilcoxon signed-rank test, p-value = 0.86), although π is significantly lower in MOX compared to SOX (two-sided Wilcoxon signed-rank test, p-value = 5.23e-4). The lower coverage of the MOX data could cause difficulties in estimating accurate gene frequencies and result in an overestimation of φ. To investigate the impact of coverage on the φ estimation, we estimated φ for SOX downsampled to the MOX coverage. We found a strong correlation between SOX φ and φ for the downsampled data, where the downsampling resulted in a slight underestimation of φ (Fig. S4C). We therefore conclude that the high values of MOX φ cannot be explained by the lower MOX coverage.

**Figure 2.**
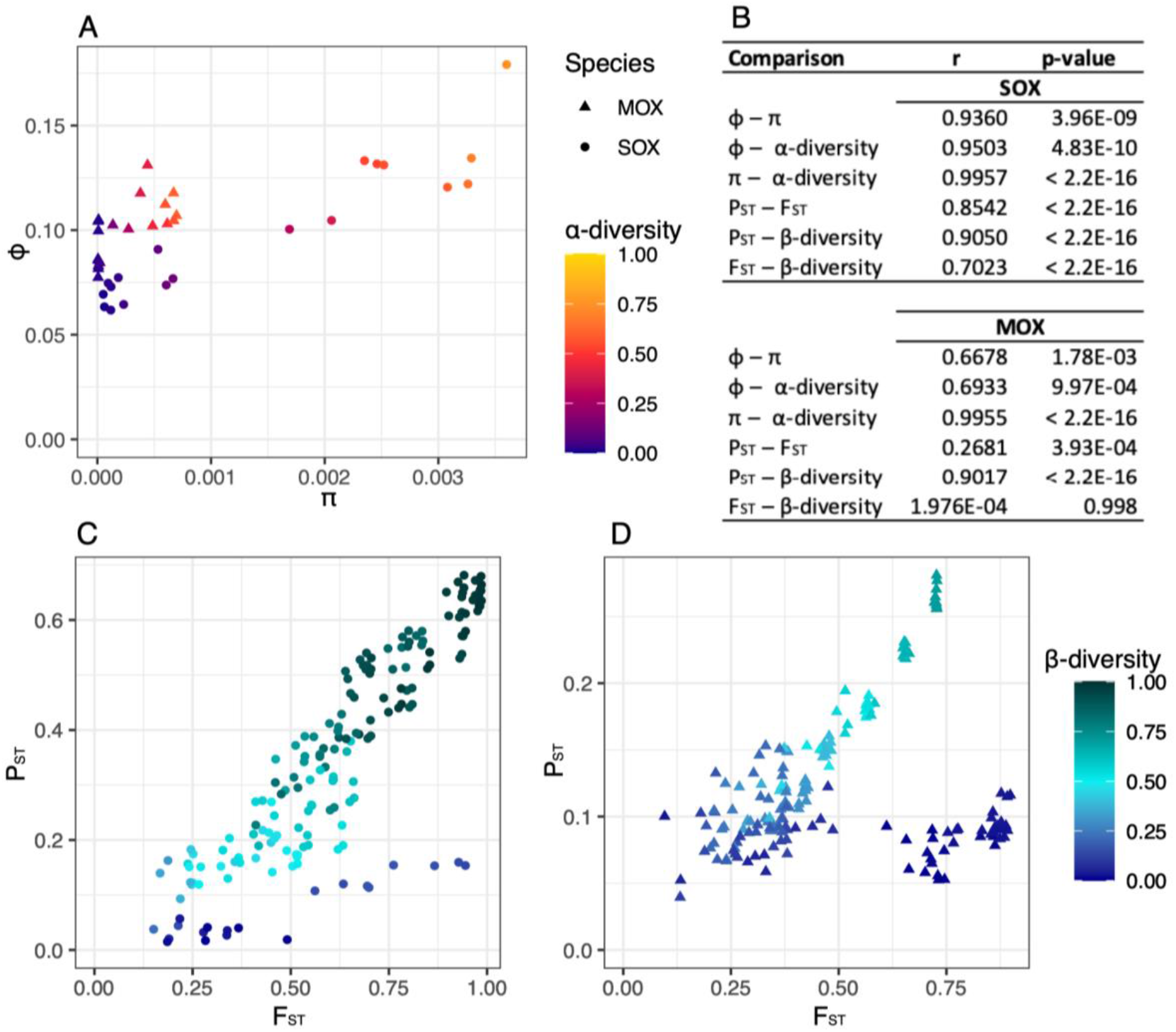
Relationships between different measures for population diversity and genetic isolation. **A)** Relationships of gene content diversity (φ) with nucleotide diversity (π) and α-diversity for SOX and MOX. **B)** Correlation coefficients and p-values for Pearson’s product-moment correlation. **C)** Relationships of pangenome fixation index (P_ST_) with fixation index (F_ST_) and β-diversity for SOX. P_ST_ (median 0.3417) is significantly lower than F_ST_ (median 0.6216) (two-sided Wilcoxon signed-rank test, p-value < 2.2e-16). **D)** Relationships of P_ST_ with F_ST_ and β-diversity for MOX. P_ST_ (median 0.1043) is significantly lower than F_ST_ (median 0.4290) (two-sided Wilcoxon signed-rank test, p-value < 2.2e-16).

To study the degree of isolation between mussels, we developed the pangenome fixation index P_ST_, which is based on the gene content diversity φ and derived in analogy to the fixation index (F_ST_) (see Methods). Small values of F_ST_ or P_ST_ indicate that the samples stem from the same population, whereas large values indicate that the samples constitute subpopulations. We found that P_ST_ measures a lower degree of isolation than F_ST_, which is particularly pronounced in MOX (Fig. 2C,D, S5). Despite P_ST_ being lower than F_ST_, both measures are correlated, where the correlation is especially strong for SOX. Furthermore, we observed that for both SOX and MOX, the pairwise correlation is highest when comparing P_ST_ and β-diversity. The β-diversity measure is based on the strain relationships and the strain distribution in the mussels (Fig. 1). Thus, differences in strain composition are even more strongly correlated to the pangenome fixation index than to the SNV-based fixation index. We have observed before that some pairwise comparisons have high F_ST_ but low β-diversity, especially for MOX (Romero Picazo et al. 2019). Notably, all SNVs are considered for the F_ST_ estimation, but not all SNVs can be linked to strains by DESMAN, which can result in discrepancies between these two measures. Here we observed that P_ST_ is low for these discrepant pairs resulting in the dark blue clouds in the bottom right part of Fig. 2C,D. Thus, P_ST_ reflects β-diversity stronger than it reflects F_ST_. We also observed that samples cluster based on these diversity measures, where the clustering based on P_ST_ is very similar to the clustering based on β-diversity for both symbionts (Fig. S4).

Taken together, we derived methods to study population diversity and genetic isolation based on gene content. We find that the measured degree of isolation is stronger when measured at the level of SNVs (i.e., F_ST_) in comparison to gene content (i.e., P_ST_), which might be due to the fact that SNVs and their frequencies can be measured with a higher accuracy. We find that genetic isolation based on gene content is highly associated with β-diversity based on strain composition. This supports that gene content variation is strongly associated with the strain relationships instead of being mobile between strains. Since β-diversity is mostly driven by differences among the strain clades rather than differences among strains within a clade, the strong correlation further suggests that gene content differences are mainly found among strain clades. We thus conclude that gene content differences between mussels are related to the strain compositions within the mussels and especially to their clade compositions.

### Strain clades are defined by gene content differences

In the previous section, we compared the gene content within and between individual mussels. Next, we aimed to resolve the accessory gene content of individual strains by identifying the presence of genes in mussels infected by a single strain. We defined strains to be dominant in a mussel when their frequency in a sample is at least 0.7 (Table S2). For a sample with a dominant strain, genes are present in a strain when their coverage is at least 50% of the median core genome coverage in that sample and we termed all these genes to be “assigned to that strain” (Fig. S6). We further merged the strain-assigned genes of all samples with the same dominant strain. Similarly, clade-assigned genes resulted from merging the gene content of strains belonging to the same clade.

For both symbionts, the majority of the accessory genes could be assigned to strains (Fig. 3); we assigned 731 genes (76% of the SOX accessory genes) to five SOX strains that are dominant in twelve mussel samples, where some genes were assigned to multiple strains (Fig. 3, Table S3A). The clades differ in the number of accessory genes, where clade S2 contains the largest number and clade S1 the smallest number of accessory genes. Additionally, S2 contains 169 genes that cannot be found in any other clade, i.e., they are clade-specific genes. In contrast, each of the other clades have at most 71 specific genes. We also identified 67 genes that were assigned to all strains, of which 18 are multi-copy genes. For MOX, we could assign 276 accessory genes (67% of the MOX accessory genes) to three different strains that were dominant in ten different samples (Fig. 3, Table S3B). Of these, 65 genes are specific to clade M1, 132 are specific to clade M2, and 77 genes were shared among all three strains, of which 34 are multi-copy. We found that the reconstructed gene content of samples with the same dominant strain overlap to a large extent, which serves as additional support for the robustness of our approach (Table S3).

**Figure 3.**
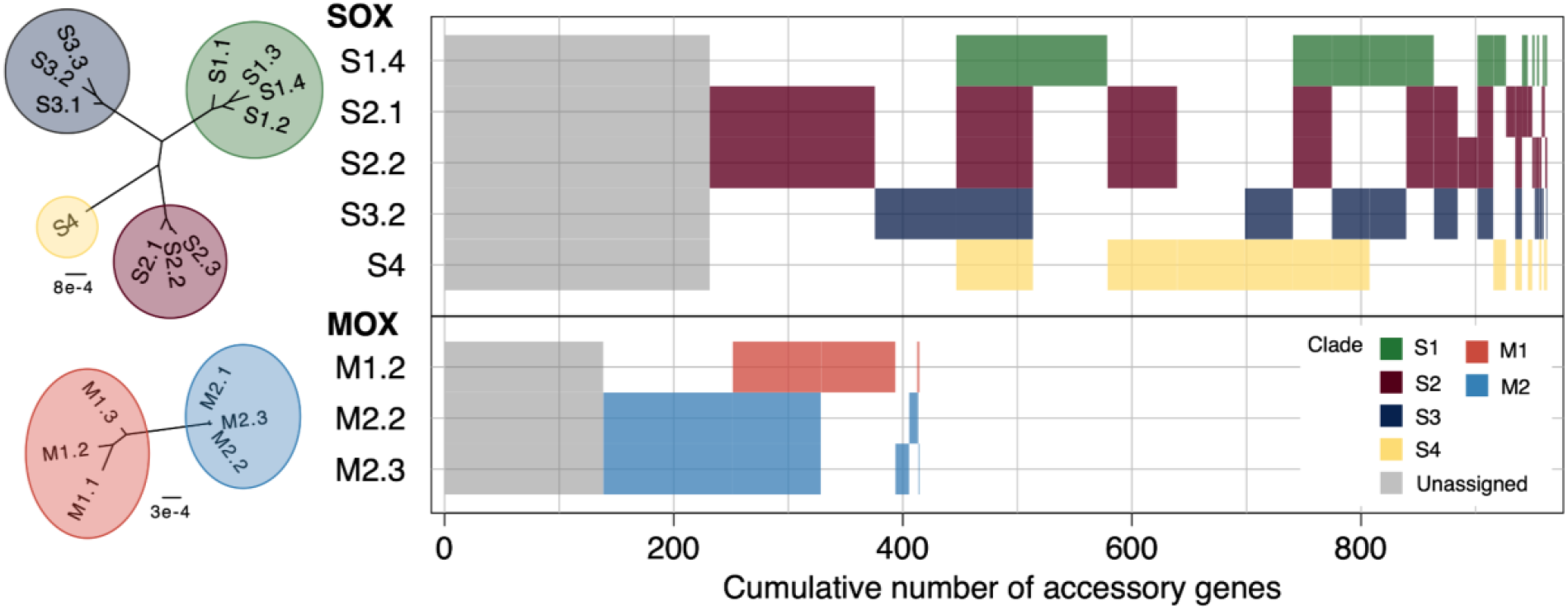
Accessory gene content of reconstructed strains in SOX and MOX. Presence-absence patterns for accessory gene clusters are shown. Gray gene clusters refer to accessory genes that could not be assigned to any strain, whereas colored gene clusters indicate the clade affiliation. Blocks of gene clusters are sorted by size. For interpretation, core strain phylogenies are shown on the left. Based on the concatenated core gene alignment, IQ-TREE 2 using the GTR+G8 model and 1000 approximate bootstrap replicates resolved these trees with all branches supported >85% (Minh et al. 2020). A network representation of the strain relationships is presented in Fig. 1 of (Romero Picazo et al. 2019).

To gain insights into the pace of gene content evolution, we estimated pairwise strain distances in terms of differentially present genes and of SNVs. We found that strain pairs generally have more SNVs compared to differentially present genes (Fig. S7, Table S4), which supports that gene content changes occur less frequently than substitutions. The positive correlation between the number of differentially present genes and SNVs supports a positive relationship between gene content changes and substitutions, which is compatible with gene content evolution by vertical inheritance.

Here, by extracting genes present in samples with a dominant strain, we could assign a large proportion of accessory genes to SOX and MOX strains. We found a high level of gene sharing among strains within the same clade, which supports that clades are characterized by gene content and that gene content evolves mainly by vertical inheritance.

### The accessory genome is less diverged than the core genome for both symbionts

To investigate the evolution of the accessory gene sequences, we identified SNVs on all the genes in the pangenomes. The number of SNVs per kilo base pair (SNVs/kbp) is higher in SOX than in MOX for all gene classes (core, single-copy accessory, and multi-copy accessory) (Table 1). For both species, the SNVs/kbp is lower in the accessory genome than in the core genome, where many of the single-copy accessory genes do not have any SNV (Fig. S8A, Table 1) - 475 genes in SOX (51%) and 298 (79%) in MOX. Furthermore, multi-copy genes have the highest SNVs/kbp (Table 1), which suggests that the divergence in multi-copy genes is overestimated due to the inclusion of paralogs.

To study the selection pressure on the pangenomes, we estimated pN/pS for the single-copy genes, where the analysis is restricted to genes with at least one SNV. In MOX, core and accessory genes show a similar pS distribution and also a similar pN/pS distribution (Fig. S8B,C). Thus, although divergence is low in MOX, the selection pressure acting on the core and accessory genome is similar. In contrast, we observed that the pN/pS distribution is shifted to lower values for the SOX core genes compared to SOX accessory genes (Table 1, Fig. S8B). This could suggest that the strength of purifying selection is higher on the core genome. However, the pS distribution is shifted to higher values for the SOX core genes compared to the accessory genes (Table 1, Fig. S8C). It has been observed that the relative rate of nonsynonymous to synonymous substitutions depends on the divergence of the analyzed species (Rocha et al. 2006; Romero Picazo et al. 2019). Indeed, we found that the joint distribution of pS and pN/pS are largely overlapping for the SOX core and accessory genome, with some accessory genes having a very low pS and a high pN/pS (Fig. S9A). We thus cannot conclude that the strength of selection is different between SOX core and accessory genes.

To rule out the possible impact of differential coverage between core and accessory genes in the pN and pS estimates, we studied the distribution of these measures by only considering SNVs found across those samples containing a dominant strain. In these samples, SNV detection should be less biased by coverage because core and accessory genes have similar coverage (Fig. S6). Indeed, we found that the distributions of pN/pS and pS estimated only from those samples are similar to the distributions previously estimated for the full dataset (Fig. S9), which suggests that variation in coverage does not explain the differences observed between the core and accessory genome.

The frequencies of SNVs/kbp presented here suggest that the accessory gene sequences are less divergent than the core genes in both symbionts. In a scenario where the accessory genes are enriched for mobile genes and have been taken up multiple times from the environmental gene pool, the accessory genome is expected to be very diverse. We thus conclude that this scenario is not realistic. In contrast, the majority of the accessory genome has probably only been acquired once or has been present in the ancestor of the population and was subsequently lost in some strains. Then, accessory genes are found at lower frequency and are thus expected to be less diverged than core genes. Furthermore, the larger number of genes without detected mutations in the accessory genome of MOX compared to SOX suggests that gene family diversification is more recent in MOX compared to SOX. We could not detect a difference in selection pressure acting on the core and accessory genomes, mainly due to the low divergence level of these genes.

### Genes in the accessory genome mostly function in genome integrity

To further investigate the origin of the accessory genes in the MOX and SOX symbiont populations, we studied the distribution of functional categories across the core and accessory genomes. Functional annotation of all genes in the pangenomes by clusters of orthologous groups (COGs) revealed that functions associated with central metabolism are overrepresented in the core genomes of both SOX and MOX pangenomes (COG categories “translation, ribosomal structure and biogenesis” and “amino-acid transport and metabolism”). In SOX, multiple additional categories related to central metabolism are overrepresented (Fig. 4A,B).

**Figure 4.**
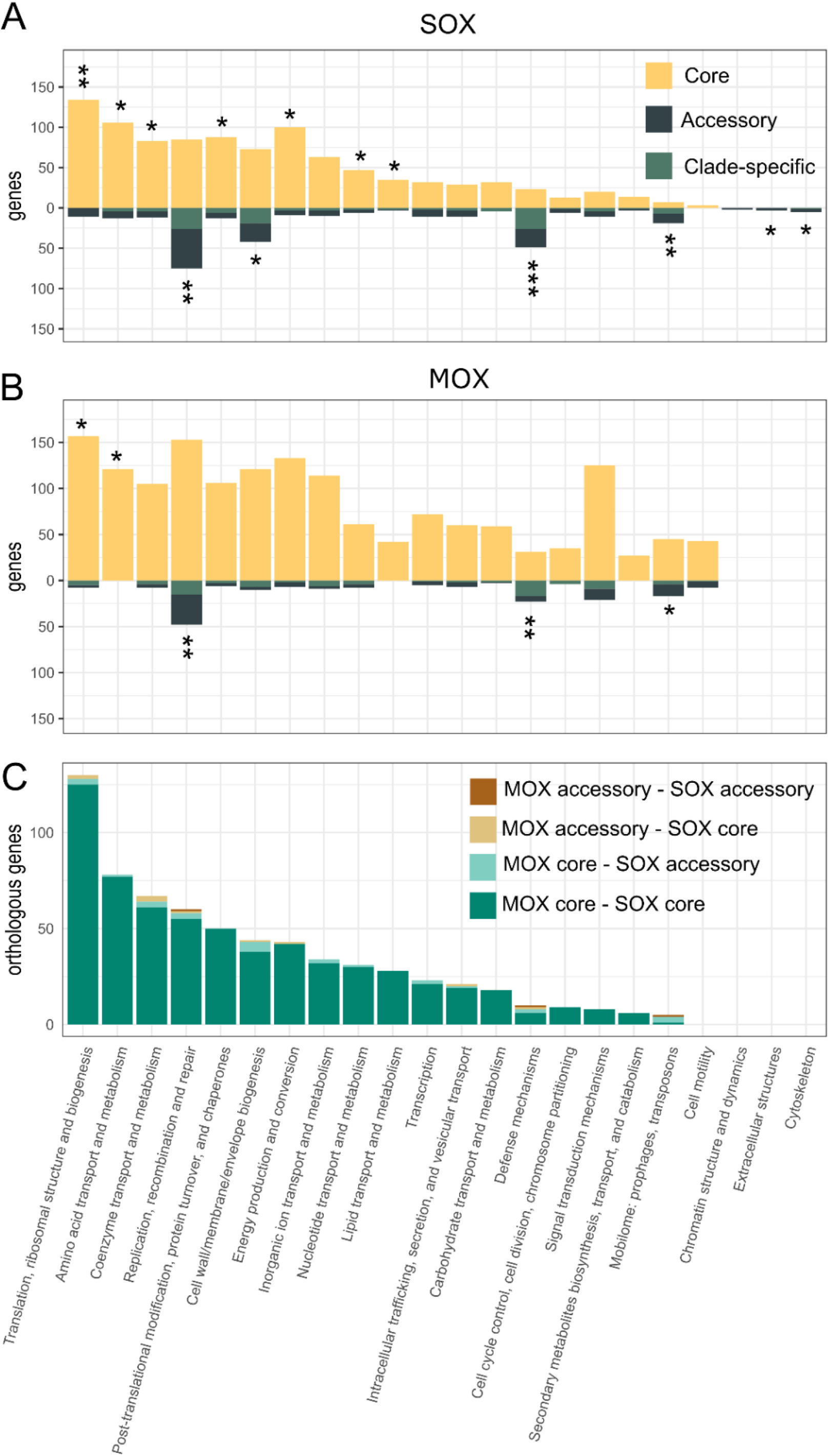
COG annotations for SOX and MOX pangenomes. Distribution of COG annotations for **A)** SOX and **B)** MOX core and accessory genomes. Stars represent the significance of Fisher’s exact test for the differential presence of a specific COG category between core and accessory genes. * - p-value < 0.05, ** - p-value < 5×10^−5^, *** p-value < 5×10^−10^ (FDR-corrected p-values). **C)** Distribution of COG categories for 760 homologous pairs between SOX and MOX pangenomes, where 51 homologs involve at least one accessory gene (functions listed in Table S7). Ten homologs are accessory in both genomes (incl. two transposases and two restriction-modification system genes). Twenty-eight pairs involve a core gene in MOX and an accessory gene in SOX (incl. 5 genes in the COG category ‘Cell wall/membrane/envelope biogenesis’, and also transposases, integrases and restriction-modification system genes). Thirteen pairs involve a SOX core and a MOX accessory gene (incl. 3 genes in the COG category ‘Coenzyme transport and metabolism’).

The accessory genomes of both populations are overrepresented in functions associated with genome integrity, namely “Mobilome: prophages, transposons”, “Defense mechanisms” and “DNA replication, recombination and repair”. The COG category “Mobilome: prophages, transposons” is more prevalent in the MOX pangenome (62 genes in MOX, 26 genes in SOX). However, SOX shows a larger proportion of this category in the accessory genome (19, 73%) than MOX (17, 27%). Investigating the functional annotation of the genes in detail, we identified 14 integrases in SOX and five in MOX. Additionally, we found 70 transposases in MOX and 20 in SOX. Of these, 16 (23%) were identified as multi-copy genes in MOX, while three (15%) were identified as multi-copy genes in SOX.

The COG category “Defense mechanisms” is overrepresented in the accessory genomes of both populations (49 accessory out of 72 genes in SOX, 68%; 23 accessory out of 54 genes in MOX, 43%). We found a high number of genes associated with restriction-modification systems: 72 in MOX, where 24 (33%) are accessory, and 81 in SOX, where 48 genes (59%) are accessory. Additionally, we found a larger repertoire of CRISPR-Cas genes in SOX (20 genes, thereof 16 accessory) in comparison to MOX (seven genes, thereof none accessory). The prevalence of defense mechanisms opens the question, whether viruses are present in the mussel environment. The prediction of phages on all metagenomic contigs revealed 51 potential viral contigs, of which there is only one high-confidence complete phage contig (termed *Gokushovirinae* sp. isolate VC_68_0, Table S5). Thus, viruses are either not highly abundant in the *B. brooksi* microbiome or could not be detected in the data analyzed here.

Several functions are differentially present between the pangenomes of the symbionts (Fig. 3A,B). For example, we found that MOX contains 51 genes related to “Cell motility”, where 43 (84%) are in the core genome, whereas only three core genes were identified in SOX for this category. Additionally, 146 genes were annotated as “Signal transduction mechanisms” in MOX (21 accessory genes, 14%), whereas only 31 genes were found in SOX (11 accessory genes, 35%). To further examine the functions that are differentially present among strains, we inspected the genes that have been assigned to strains (Table S6). We observed that most of the strain-specific genes in SOX belong to the categories “Defense mechanisms” and “Replication, recombination and repair”, followed by “Cell wall/membrane/envelope biogenesis” category. Notably the COG category “Cell wall/membrane/envelope biogenesis” also includes the toxin-related genes that have been described previously to be variable across strains (Sayavedra et al. 2015). In MOX, the categories “Defense mechanisms”, “Replication, recombination and repair”, and “Cell wall/membrane/envelope biogenesis” also differ between strains and, additionally, MOX strains differ in genes involved in “Signal transduction mechanisms”. For SOX and MOX, categories related to metabolism are rarely found to be strain-specific. The differences among the strain-specific functional categories suggest that the different strains differ in the repertoire of genes that are involved in interactions between organisms, including interactions with the mussel host and with other bacteria or mobile elements.

We conclude that in both symbiont pangenomes, the accessory genomes comprise mostly gene functions related to genome integrity. The high prevalence of transposases and restriction-modification systems suggests that genome rearrangements contribute to gene content variation among strains. Additionally, we found that MOX has a larger repertoire of mobilome-related genes and contains more genes related to cell motility and signal transduction.

### No evidence for horizontal gene transfer between both symbiont species

For endosymbiotic bacteria, the concept of an “intracellular arena” posits that the host cell serves as an arena for bacteria, where symbionts can acquire putatively beneficial genes from a niche-specific gene pool (Bordenstein & Reznikoff 2005; Newton & Bordenstein 2011). To identify mobile elements with the potential to transfer genes in that environment, we studied the diversity of transposases in the pangenomes. Transposases can duplicate within or between genomes, where recent duplications will show a low diversity between both transposase copies. Here, we observed a low genetic diversity only among transposases within the same species, whereas the minimum divergence between species is 0.50 substitutions per site (Fig. S10). Thus, recent transposase duplication events have occurred only within species and not between species, which leads to the conclusion that there is no evidence for recent transfer of transposases between both symbiont species.

In addition, we aimed to identify possible HGT events between SOX and MOX symbionts. To this end, we inferred 760 homologous protein pairs between the two symbionts. Of these, 709 (93%) comprise core genes from both species, where the majority encode for central metabolism functions (Fig. 4C). The homologs that are core genes of both species are presumably related by common ancestry, i.e., they are orthologs. In addition, 51 homologs involve at least one accessory gene (Table S7, Fig. 4C). These are candidates for a recent horizontal transfer between symbionts, where a transfer would result in a high sequence similarity between the homologs. However, the sequence identity between homologous pairs does not reach high values (less than 86%, Fig. 5). In addition, the distributions of sequence identity of core pairs and of pairs involving at least one accessory gene are not significantly different (Wilcoxon rank sum test p-value= 0.3502; Fig. 5). We conclude that the homologous pairs involving an accessory gene are indeed orthologs, which were differentially lost in one or several strains, rather than being acquired recently by HGT between the symbionts.

**Figure 5.**
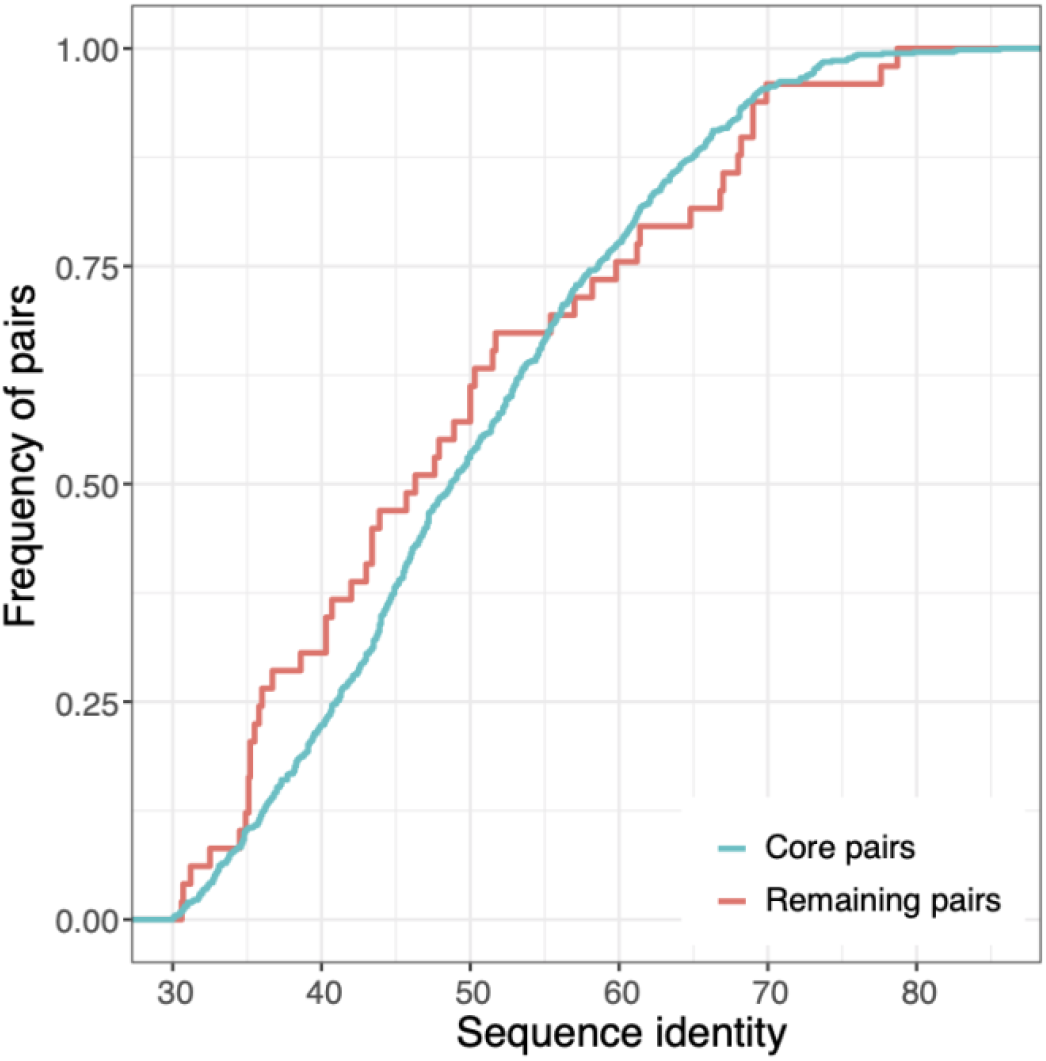
Empirical cumulative distribution of sequence identity between orthologous pairs. Core pairs sequence identity: median 48.7, maximum 85.6; remaining pairs (i.e., containing at least one accessory gene) sequence identity: median 46.3, maximum 78.7.

Note that our approach might not be able to detect very recent transfers between SOX and MOX where both homologs are present in all of the samples. This limitation exists because a very recent between-species transfer within a single mussel may lead to discontinuous assemblies and exclusion of the transferred gene from both SOX and MOX contigs. However, transferred genes that accumulated mutations in the donor or recipient should be distinguishable during the assembly. Our approach is expected to perform well in detecting HGT events, where homologs are differentially present in the different mussels or where homologs accumulated more than 5% nucleotide divergence. That said, we did not find any gene cluster (i.e., a group of genes with less than 5% nucleotide divergence) that is present in both pangenomes and we only found homologs between SOX and MOX with less than 86% amino acid identity, which are presumably orthologs. Consequently, we conclude that HGT between the symbiont species is rare.

## Discussion

Here we used high-resolution metagenomics to examine the gene content of two co-occurring symbiont species that inhabit mussels from a single geographical site. We reconstructed the population pangenomes of the sampling sites by applying a novel approach that links accessory genes to core genes across the chromosomal contigs of multiple samples. The pangenomes reconstructed here reflect the pangenomes of the population at the sampling site (i.e., not that of the entire species). The site pangenome is expected to be smaller compared to the species pangenome, since only populations from the same niche are included and no difference in niche-specific genes is expected. We thus expect that the SOX and MOX species pangenomes are larger with a larger proportion of accessory genomes than the site pangenomes reported here.

The reconstruction of pangenomes from short-read metagenomic sequencing data is a challenging task. When isolate genomes are available, pangenome analysis can include the mapping of metagenomes to infer the ecology of the species (Delmont & Eren 2018; Utter et al. 2020). For species that cannot be cultivated, however, only metagenome-assembled genomes (MAGs) might be available for a pangenome analysis. Although MAGs might miss genomic regions compared to isolates (Meziti et al. 2021; Nelson et al. 2020), pan-genomics based on MAGs is a widely used approach for studying the evolution of bacteria that are difficult to cultivate; recent analyses include *Wolbachia* sequenced with their hosts (Scholz et al. 2020), *Sulfurovum* from deep-sea hydrothermal vents (Moulana et al. 2020), Thaumarchaeota from river sediments (Sheridan et al. 2020), and Chlamydiae present in various habitats covered by the Earth Microbiome initiative (Köstlbacher et al. 2021). It is particularly difficult to study genome rearrangements and to infer unlinked mobile genetic elements such as plasmids from short-read metagenomes (Maguire et al. 2020; Nelson et al. 2020). Additional approaches such as isolation and sequencing, methylation patterns from long reads (Beaulaurier et al. 2018), or linking DNA by Hi-C (Yaffe & Relman 2020) would be necessary to resolve mobile genetic elements with confidence and to link them to their host. The inferred pangenomes reported here thus comprise the core and accessory genes located on the symbiont chromosomes. We use several steps to improve the accuracy of the pangenome reconstruction from metagenomes (Fig. 1). First, we employ co-abundance binning to reconstruct core genomes and subsequently a network approach to include all accessory genes that are linked on contigs to core genes or accessory genes from other samples. Second, we estimate the coverages for all genes in the pangenomes in each sample, also including genes that have not been reconstructed on the contigs of that sample. Third, strain content is highly variable across samples, and we observed several samples that were dominated by one strain only. These strains show very good assembly statistics (e.g., SOX N50 above 30,000, Table S2) and have been used to infer the strains’ gene contents. We thus conclude that the reconstructed MAGs and the reconstructed pangenomes are of high quality. Finally, our analysis focuses especially on the genes that can be assigned to strain clades; these have been maintained over long time scales and might have an evolutionary relevance. However, we might miss low-frequency strains in our analysis and accessory genes that are only present in samples with a high strain diversity might be missing from the assemblies. However these missing genes are not yet fixed in a strain clade and might thus not be relevant for adaptation. We thus conclude that the potentially missing genes are transient and belong to the cloud genome, i.e. they are rare or nearly unique genes (Koonin & Wolf 2008).

Whereas the SOX pangenome has less genes than that of MOX, the former has a higher fraction of accessory genes. The large SOX accessory genome is consistent with the recent finding that gene content variation among coexisting thiotrophic bacteria is common (Ansorge et al. 2020). We find that functions associated with genome integrity are consistently present in the accessory genomes of both species. Among them are genes encoding for defense mechanisms, in particular, genes related to restriction-modification systems. In addition to defense, restriction-modification systems can also function as mobile genetic elements. They have, for example, been shown to be involved in genome rearrangements in termite gut symbionts and in gene birth and death in the human gut bacterium *Helicobacter pylori* (Furuta et al. 2011; Zheng et al. 2016).

We find that mobile genetic elements, such as transposons, are more prevalent in MOX, which also has a high proportion of them in the core genome. Furthermore, MOX has a higher fraction of genes related to cell motility and signal transduction, which can be found in the core and accessory genome. Notably, both these functional categories have been found to be underrepresented in intracellular compared to free-living bacteria (Merhej et al. 2009; Lo et al. 2016); thus, they are more relevant for a free-living lifestyle. This suggests that MOX still pursues an active free-living life stage or that the association of MOX with *Bathymodiolus* is recent and ancient genes can still be found in the genome. Mussel phylogenies indeed support that the association with MOX is younger than that with SOX, where the clade comprising *B. brooksi* evolved from an ancestor with only the SOX symbiont about 10 million years ago (Lorion et al. 2013). Nevertheless, co-speciation of hosts and symbionts is rare in that system (Won et al. 2008) and it can thus not be ruled out that SOX and MOX symbiont populations have been replaced multiple times during mussel evolution.

Differences in pangenome size can also be caused by different rates of HGT. Nevertheless, we conclude that HGT is rare between species and between strain clades of both species (Fig. 6). This conclusion is supported by the observations (i) that gene content differences between mussel individuals reflect the differences in strain composition, (ii) that the accessory genome is less diverged than the core genome for both symbionts, and (iii) that the homologs between SOX and MOX are not highly similar. Observations (i) and (ii) support that accessory genes have been acquired at most once from the environmental gene pool and then evolved by descent with modification within the symbiont lineages. Gene loss or recombination within the strains might also contribute to accessory gene evolution, whereas multiple transfer from the environmental pool or horizontal gene transfer between different strain clades is rare. Since additionally the reconstruction of homologs between SOX and MOX also did not reveal signals of recent gene transfer between the species (observation iii), we conclude that also HGT between the two species is rare. This contrasts their common ecology and their potential ability to access the habitat-specific gene pool as described for other species (Bordenstein & Reznikoff 2005; Newton & Bordenstein 2011; Polz et al. 2013). Notably, in SOX from different *Bathymodiolus* host species, a higher fraction of genes that potentially originated by HGT has been previously found (Sayavedra et al. 2015). Considering that our analysis is restricted to events within the SOX population at a particular site, this discrepancy may stem from differences in the sampling distribution of the compared strains. Furthermore, host-symbiont interactions can differ on the molecular level between different *Bathymodiolus* species and their symbionts (Ponnudurai et al. 2017, 2020), which might also result in differences in the evolution between symbionts of different *Bathymodiolus* species.

**Figure 6.**
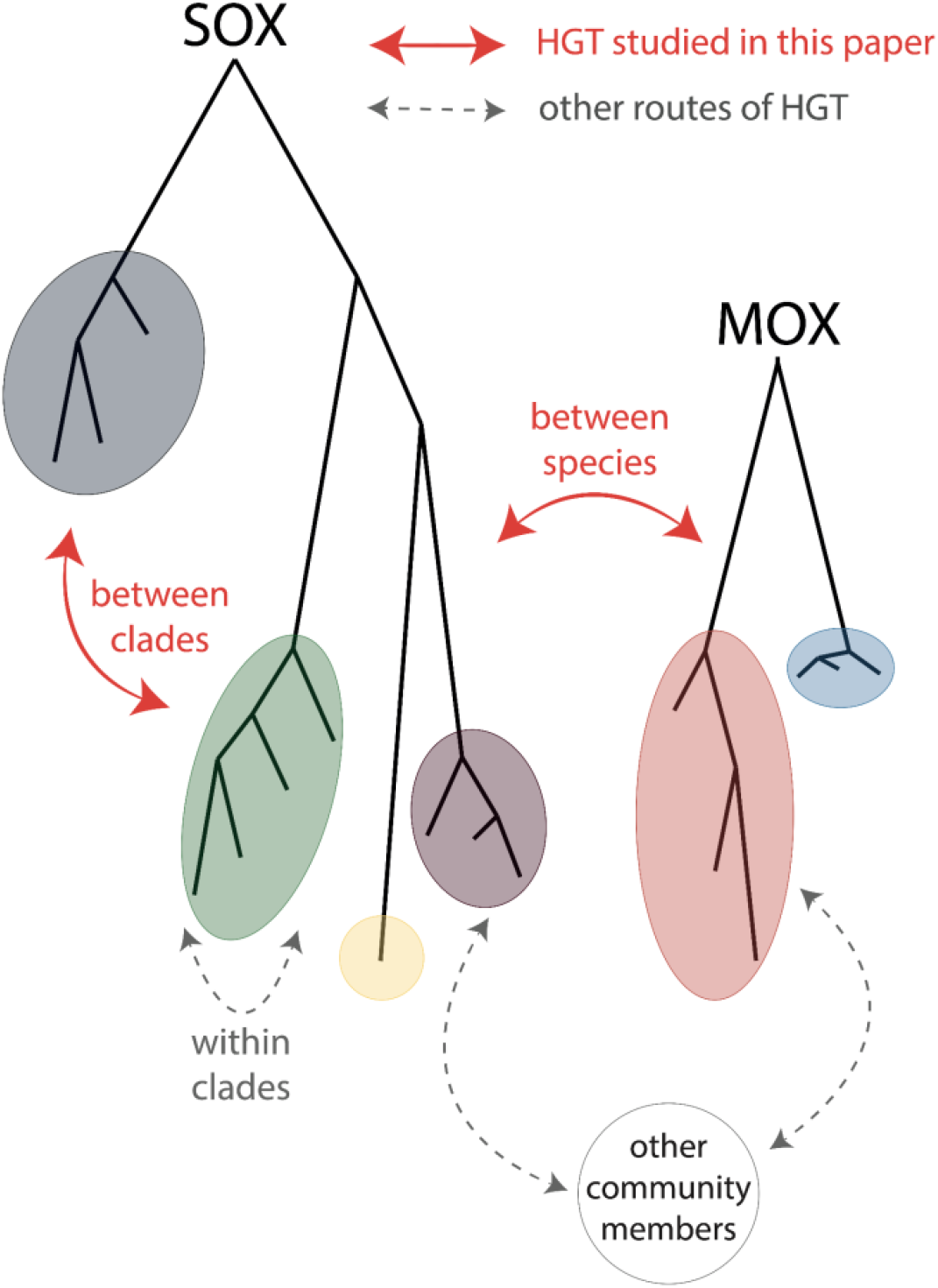
Routes of HGT studied here.

Notably, our conclusion that HGT is rare for SOX and MOX contrasts what is known for most other bacterial species, where HGT is a major evolutionary driver (Treangen & Rocha 2011; Brockhurst et al. 2019). The rarity of HGT in this environment might be explained by the absence of DNA transfer mechanisms in the symbionts or by environmental properties. Regarding the latter, the intracellular environment may interfere with mechanisms that rely on the transfer of free DNA such as natural transformation, since the DNA could be quickly degraded by the prevalent mussel digestive enzymes (Ponnudurai et al. 2017). Likely HGT mechanisms in such an environment are such where the DNA is transferred in a packaged manner (such as in phages, gene transfer agents, or outer membrane vesicles) or transferred in direct contact between donor and recipient (as in conjugation). However, these mechanisms are most likely to transfer genes between symbionts within a single bacteriocyte and only a few symbionts are generally harbored within a single bacteriocyte (Dubilier et al. 1998). We thus conclude that potential gene transfer events rarely establish in the population and that the host association results in genetically isolated subpopulations where HGT is limited. This is consistent with the observation that endosymbionts are rarely connected in gene transfer networks (Popa et al. 2011).

Symbioses are traditionally distinguished by being open (symbionts are environmentally acquired, resulting in frequent HGT and average or high GC content), closed (symbionts are parentally transmitted, resulting in the absence of HGT, low GC content, and genome degradation), or mixed (symbionts are mainly parentally transmitted, but occasional environmental acquisition occurs) (Perreau & Moran 2022). However, the *Bathymodiolus* symbionts are environmentally transmitted, do not engage frequently in HGT, and have an average GC content of 38%. Thus this symbiosis does not fit into the traditional division. In invertebrates, the innate immune system has an important role in establishing symbioses resulting in highly specific symbiont acquisition by the host from the environment (Nyholm & Graf 2012). We conclude that these symbioses are not open, but rather restricted, thus we term them *narrow* symbioses. In narrow symbioses, the tight host association leads to genetically isolated subpopulations with low frequencies of gene transfer within the host environment, where low rates of recombination can rescue the genomes from extensive degradation (Russell et al. 2020). Pangenome evolution differs substantially between open and narrow symbioses, where gene content evolution of the latter symbionts is mainly driven by differential gene loss and HGT happens only occasionally.

## Materials and Methods

### Collection, sequencing, and core genome reconstruction

Details of the sample processing and sequencing have been described previously (Romero Picazo et al. 2019). In brief, 23 *Bathymodiolus brooksi* mussels were collected from a cold seep location at the northern Gulf of Mexico and mussel gill homogenate has been sequenced using Illumina HiSeq2500 (250 bp paired-end reads with a median insert size of 400 bp, raw reads BioProject PRJNA508280). The core genomes are reconstructed as described previously (see also Fig. 1). In brief, individual samples were assembled with metaSPAdes (Nurk et al. 2017) and predicted genes from all samples were clustered with identity of 95% into gene clusters that represent the non-redundant gene catalog. Next, we estimated the gene abundances in each sample and performed co-abundance gene segregation by using a canopy clustering algorithm (Nielsen et al. 2014), which groups gene clusters into bins that covary in their abundances across the different samples. Two metagenomic species (MGS) were classified as the SOX and MOX core genome, respectively, and lowly and highly covered genes were further filtered. We additionally identified a third MGS (MGS3), which is present in a single sample in low abundance and for which no taxonomy could be inferred. After discarding samples with high variance in symbiont marker gene coverages and with low coverage after binning, 19 mussel samples remained as the basis for the following analysis. In addition to the analysis presented in (Romero Picazo et al. 2019), singletons that originated from discarded samples were removed from the core genomes. This results in a SOX core genome of 1,408 genes and a MOX core genome of 2,443 genes.

### Pangenome reconstruction

Differences in strain composition generate different assembly fragmentation patterns across samples, where genome regions that are present only in specific strains tend to result in isolated contigs. Here, we restore the linkage between contigs that belong to the same species and identify, from all the genes present in the catalog, the accessory and multi-copy genes that belong to SOX and MOX symbiont pangenomes. Our approach is based on the non-redundant gene catalog, which contains gene clusters present across samples with at least 95% of sequence identity (Fig. 1).

We used a network traversal approach, where the genes that are part of core gene clusters were used as initial seeds. First, we located the seed genes on the contigs of the 19 samples. The first layer of the pangenome contains all additional gene clusters having gene members that are found on any of those contigs. Next, we used the genes that are part of these newly added gene clusters as seeds to expand the network. The extension of the network continues in an iterative manner until no additional genes can be added to the pangenome (Table S1). Microbial genomes contain multi-copy genes such as transposons. The use of multi-copy genes as seeds would potentially cause the spurious linkage of genome fragments. In order to avoid such artifacts, we only considered those genes as seeds that originate from clusters with sequence identity of at least 0.95 and that contain at most one gene from each sample. Nevertheless, as multi-copy genes can naturally be found in bacterial genomes, they were included in the pangenomes. We discarded a singleton gene from the SOX core genome connecting to the MOX pangenome. The link between the two pangenomes occurred in the second layer of the network, when two contigs containing each two genes in each pangenome were connected by another contig containing two genes. Although the SOX singleton certainly has coverage corresponding to a core gene, it only is present in one sample and has no functional annotation. Therefore, we suspect that the singleton has been misassigned as a SOX core gene and decided to not include it in the analysis. To avoid misassignment of genes in the MOX pangenome, we additionally discarded the two genes belonging to the contig connecting both pangenomes as well.

The quality of the pangenomes has been assessed by studying the distribution of two different gene cluster features across the pangenome layers, the cluster size and the cluster sequence identity (Table S1). For clusters added to layers of the pangenome, the gene cluster size is below 19 (the number of samples). This is expected for gene clusters that are not present in every strain and supports their status as accessory. Additionally, the median sequence identity of the clusters is close to one (>0.99), which supports that the gene clusters added to the pangenomes are not affected by contamination, i.e., genes from a different MGS. We denote the additional gene clusters added to the network as “accessory” genes, where genes with a higher coverage than the maximum core gene coverage in any sample are denoted as “multi-copy accessory”, in contrast to “single-copy accessory”. In addition to the core genes, we observed genes showing a similar coverage to the median core genome coverage across samples (median sample coverage ± 0.15 x median core sample coverage for all samples, 114 SOX genes, 9 MOX genes). These genes were added to the core genome for all the presented analyses.

The contigs connected in the network of each species correspond to the metagenome-assembled genomes (MAG) of each sample. We assessed the degree of completeness and contamination for MOX and SOX MAGs using the set of Gammaproteobacteria marker genes and the “taxonomy_wf” workflow from CheckM v1.0.18 (Table S2) (Parks et al. 2015).

### Measures of population diversity and diversification

Single nucleotide variants (SNVs) were detected on the non-redundant gene catalog as described in (Romero Picazo et al. 2019). Briefly, the reads in each sample were first downsampled to the minimum number of reads per sample and mapped to the gene catalog using bwa mem (Li 2013). For the analysis of the SOX pangenome, the reads of each sample were subsampled to the smallest median core coverage for SOX, i.e., when the smallest SOX core coverage is X and the sample has a SOX core coverage of Y, then all the reads were subsampled to a proportion of X/Y. For the analysis of the MOX pangenome, the subsampling was repeated based on the MOX core coverages. This normalization does not account for coverage differences due to different accessory gene abundance within a sample. LoFreq was used for probabilistic realignment and variant calling of each sample independently (Wilm et al. 2012) and the detected SNVs have been hard filtered according to GATK best practices.

Strain reconstruction for the core genomes has been performed with DESMAN as described before (Quince et al. 2017; Romero Picazo et al. 2019). In brief, this method uses the SNV frequency covariation across samples to assign the SNV states to a specific genotype. DESMAN converged with a posterior mean deviance lower than 5% for 11 SOX strains and 6 MOX strains.

The SNV data is used for calculating intra-sample and inter-sample nucleotide diversity (π) and F_ST_ as described previously (Romero Picazo et al. 2019; Schloissnig et al. 2013). π estimates the average frequency of nucleotide differences over all pairs in a sample. Small F_ST_ values indicate that the samples stem from the same population, i.e., their SNV states and frequencies are similar. In contrast, large F_ST_ values indicate that the samples constitute subpopulations, i.e., the samples differ in the frequencies of their SNV states.

To study the microbial community composition, strain frequencies and relatedness are used for calculating α- and β-diversity as described previously (Romero Picazo et al. 2019). In brief, we estimated α-diversity using phylogeny species evenness (PSE) (Helmus et al. 2007) implemented in the R package ‘Picante’ (Kembel et al. 2010). β-diversity was estimated using the weighted Unifrac distance, which quantifies differences in strain community composition between two samples and accounts for phylogenetic relationships (Chen 2018).

To estimate the degree of genetic isolation based on gene content, we have derived the intra- and inter-sample gene diversity measures. We here define the gene diversity *φ* which estimates the average frequency of gene content differences analogous to the nucleotide diversity π. For frequencies estimated from metagenomes, it is estimated as: 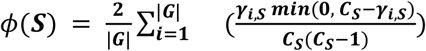 where *G* is the number of genes in the pangenome, *γ*_*i,S*_ is the coverage of gene *i* in sample *S* (measured as mean coverage across gene positions) and *C*_*s*_ is the median coverage of core genes in sample *S*. Note that the minimum ensures that the difference between accessory gene coverage and median core coverage cannot be negative. Note that this definition differs from the previously defined measure of genome fluidity, that is based on the ratio of unique gene families to the sum of gene families in pairs of genomes (Kislyuk et al. 2011). In contrast, *φ* as defined here gives an estimation for the average proportion of gene differences based on all genes in the pangenome, i.e., shared genes are only counted once.

The inter-sample gene diversity is then estimated as: 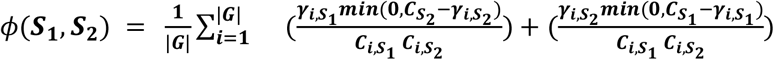, where *S*_*1*_ and *S*_*2*_ correspond to the two samples compared. Finally, analogous to F_ST_, we define the pangenome fixation index P_ST_ which measures the genetic differentiation based on the gene diversity present within and between populations: 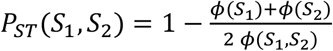. Small P values indicate that both samples have similar gene content and gene frequencies, whereas large P_ST_ values indicate that the samples differ in their gene frequencies.

pN/pS is a variant of dN/dS and is estimated based on intra-species SNVs as described previously (Romero Picazo et al. 2019). In brief, the observed ratio of nonsynonymous to synonymous mutations based on biallelic SNVs is divided by the expected ratio of nonsynonymous to synonymous mutations based on all possible mutations in each of the codons. pN/pS was estimated individually for each of the genes in the two symbiont species as well as for the core and accessory genome for each symbiont.

The scripts to calculate these statistics have been deposited at https://github.com/deropi/BathyBrooksiSymbionts.

### Reconstruction of strain gene content

To assign the accessory genes to particular strains, we first identified samples with a dominant strain, i.e. having a frequency of at least 0.7. In these samples, accessory genes with a coverage of at least 50% of the median core coverage were assigned to the dominant strain. Strain-assigned genomes were reconstructed by merging all the genes assigned to the same strain across samples. Clade-assigned genomes were inferred by merging all the genes assigned to strains of a particular clade.

### Functional annotation and overrepresentation analysis

Functional annotation by Clusters of Orthologous Groups (COGs) was determined by eggNOG-mapper v2 (Huerta-Cepas et al. 2019). Genes with multiple COGs contribute to the counts of each of the COGs. The overrepresentation of gene categories in either core or accessory genomes was done by performing multiple Fisher’s exact tests for each COG category with FDR p-value correction.

Transposase genes were identified as genes containing the substring ‘Transposase’ or ‘transposase’ and integrase genes were identified as genes containing the substring ‘Integrase’ or ‘integrase’. Additionally, we used HMMer (Eddy 2011, 2019) to screen for functional domains with hmmsearch (e-value <1e-4) against Pfam (Mistry et al. 2021) that are related to restriction-modification and CRISPR-Cas. The list of Pfam accessions used is provided in Table S8, where restriction-modification profiles have been taken from (Croucher et al. 2014) and CRISPR-Cas profiles were directly extracted by looking for the keyword ‘CRISPR-Cas’ in the Pfam portal (http://pfam.xfam.org/search).

### Phage identification

All contigs of the 19 samples were screened for viral sequences in two steps. First, contigs with VirSorter categories 1, 2, and 4 were retained (Roux et al. 2015). Second, genes on those contigs were searched with hmmscan (e-value <1e-5) (Eddy 2011, 2019) against pVOGs (Grazziotin et al. 2017) and contigs were considered as viral, if they have at least three pVOGs, where at least two pVOGs have a viral quotient of at least 0.8. Phage contigs were clustered with public phages using vConTACT2 (Jang et al. 2019). Average amino acid identity (AAI) has been calculated with CompareM (Parks 2020).

### Transposase genetic distance estimation

To estimate the degree of divergence among transposases in the population, pairwise alignments between all pairs of transposases were reconstructed with MAFFT auto mode (Katoh & Standley 2013) and pairwise distances are estimated with the K80 model implemented in the R package ape (Paradis & Schliep 2019).

### Homologous proteins between SOX and MOX

To identify homologous proteins, we extracted reciprocal best blast (Altschul et al. 1990) hits between the translated genes in the SOX and MOX pangenomes. Then, full-length protein sequence alignments were reconstructed with the Needleman-Wunsch algorithm implemented in EMBOSS (Needleman & Wunsch 1970; Rice et al. 2000), and pairs of genes with pairwise identities of at least 30% were assigned as homologous.

## Acknowledgements

We like to thank Nicole Dubilier for the collection of the original data and valuable discussions on the manuscript. We thank Marina Khachatutyan for comments on the manuscript and Franz Baumdicker for discussions on gene content diversity statistics. This work was supported by the CRC1182 Origin and Function of Metaorganisms.

## Data Availability Statement

The metagenome-assembled genomes (MAGs) of both symbionts are available at NCBI under the Bioproject ID PRJNA508280 with BioSample IDs SAMN21876924 - SAMN21876961. The genome of the virus genome Gokushovirinae sp. isolate VC_68_0 is available with the GenBank accession OL437471.1. The protein sequences for both symbiont pangenomes have been deposited in the GitHub repository https://github.com/deropi/BathyBrooksiSymbionts.

