## Supplementary figures and tables S1, S3, S4 for "Pangenome evolution in environmentally transmitted symbionts of deep-sea mussels is governed by vertical inheritance"

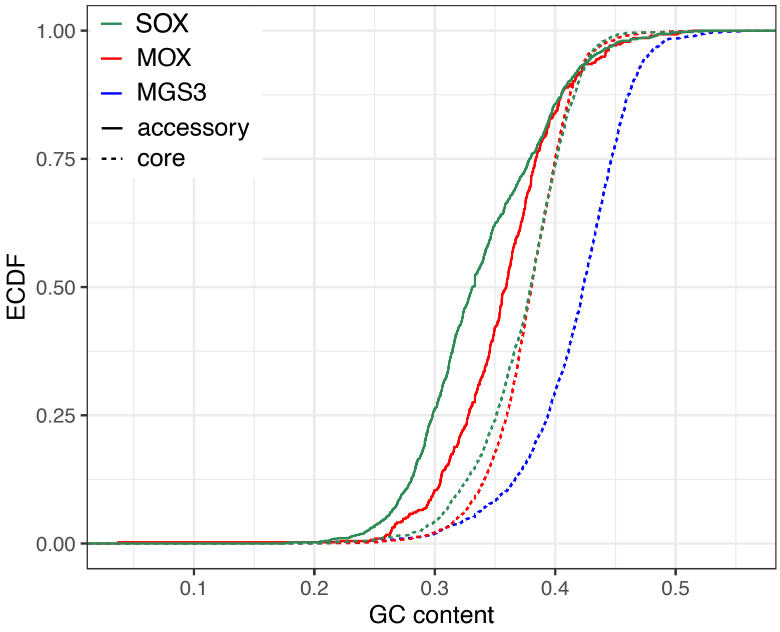


**Figure S1. Empirical cumulative distribution function (ECDF) of GC content for core and accessory genomes in SOX and MOX and for MGS3 core genome**. The accessory genomes include single-copy and multi-copy accessory genes. Median (IQR) GC content per gene for MGS3: 0.423 (0.0525), values for the symbionts are in Table 1. The GC content distributions are significantly different between the SOX and MOX core genes (Wilcoxon rank sum test, p-value = 0.03541). The GC content distribution of the SOX core genes is significantly different from the GC content distribution of the SOX accessory genes (Wilcoxon rank sum test, p-value < $2.2 \times 10^{-16}$) and the GC content distribution of the MOX core genes is significantly different from the GC content distribution of the MOX accessory genes (Wilcoxon rank sum test, p-value < $2.2 \times 10^{-16}$).

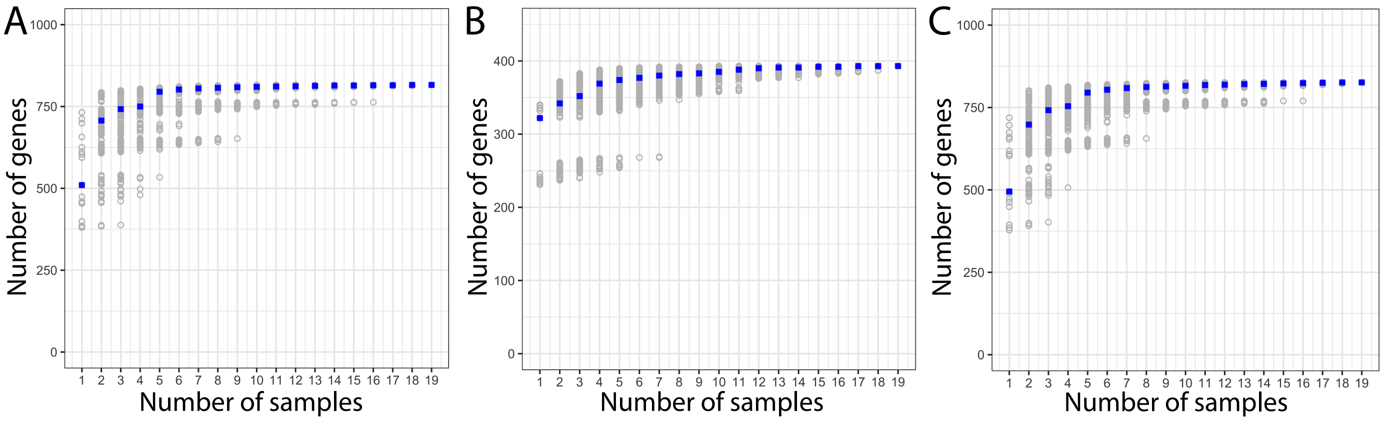


**Figure S2. Rarefaction analysis of accessory genome reconstruction.** Number of accessory genes detected in the **A)** SOX and **B)** MOX pangenome per added sample. For a sample set, a gene is denoted as detected if its coverage is at least 5% of the median coverage of the core genes in at least one sample. 1,000 random sample orders were generated and the accessory genome size (i.e., number of detected genes) is estimated; each gray dot corresponds to an individual resample; the median (blue squares) denotes the number of detected genes for each number of added strains. We observed that 95% of the total accessory genome is detected when adding five samples for both bacterial species. **C)** Rarefaction analysis of accessory genome for downsampled SOX data**.** Number of accessory genes detected in the SOX pangenome per added sample. The read data was downsampled to the median MOX core genome coverage (i.e., to a coverage of 36). A gene is denoted as detected if its coverage is at least 1.8 (i.e., 5% the median coverage of the core genes). 1,000 random sample orders were generated and the number of detected genes is estimated; each gray dot corresponds to an individual resample; blue squares show the median. We observed that 95% of the total accessory genome is detected when adding five samples.

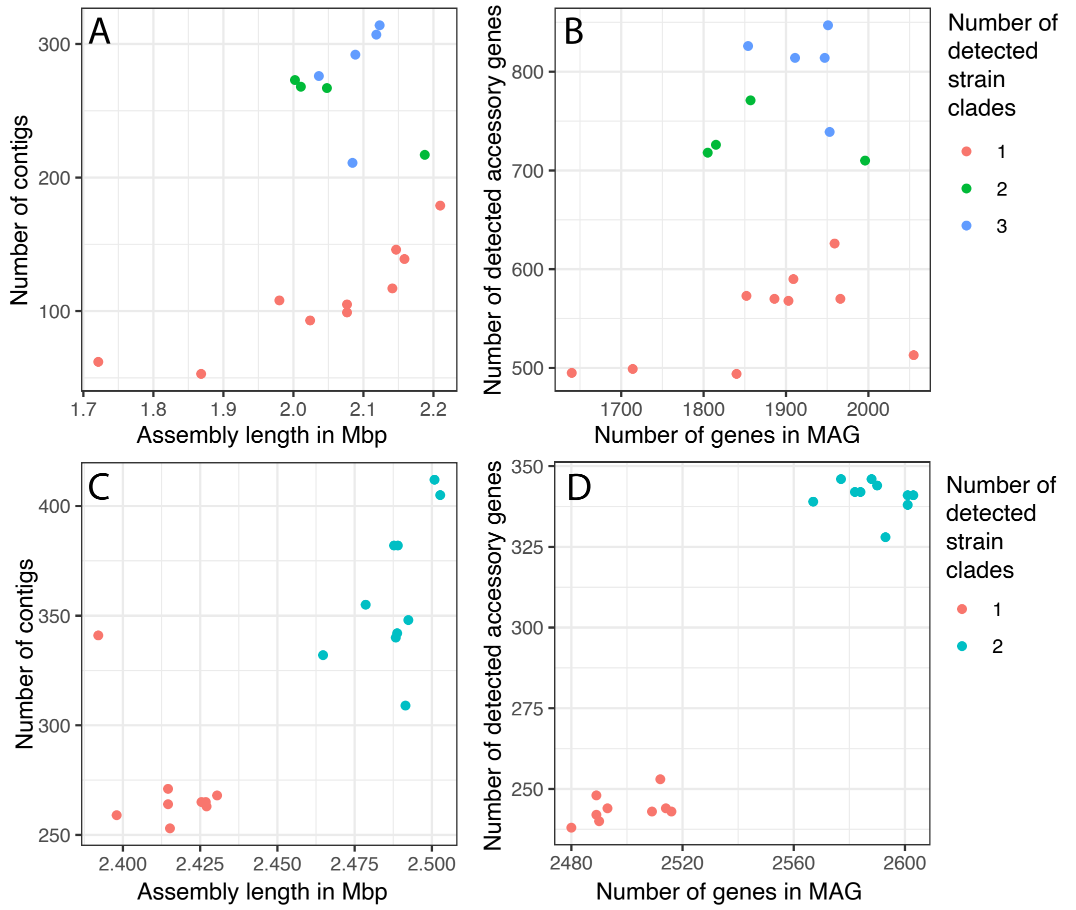


**Figure S3. Assembly quality for samples with different numbers of strain clades for A,B) SOX and C,D) MOX.** Here , a strain clade is considered detect in a sample if its frequency is at least 5%. The numbers for each sample can be found in Table S2. We observe that samples with a higher strain diversity are fragmented into a larger number of contigs. For MOX, a higher number of contigs also results in longer assembly lengths, as expected. MOX MAGs with a higher strain diversity also contain more genes and a higher number of accessory genes is detected in the sample. In contrast, in SOX the assembly length does not increase with the number of contigs and the number of genes in SOX MAGs does not increase with the number of clades. This suggests that the contig fragmentation leads to missing regions in the MAG. Nevertheless, the number of detected accessory genes increases with the strain diversity as expected.

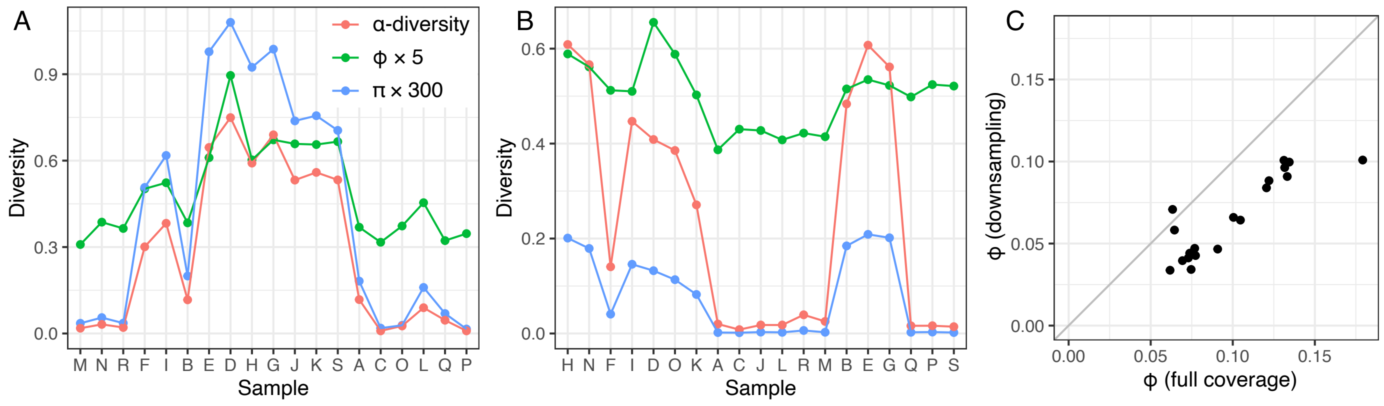


**Figure S4. Intra-sample diversity. A, B)** Nucleotide diversity (π), gene content diversity (ϕ), and α-diversity for **A)** SOX and **B)** MOX. Scaling factors are applied to π and ϕ for a better comparability of the measures. Samples are ordered according to Fig. S5. Dots are connected for visual reasons only. **C)** Effect of downsampling of SOX coverage to MOX coverage on ϕ estimation, $r^2$=0.77.

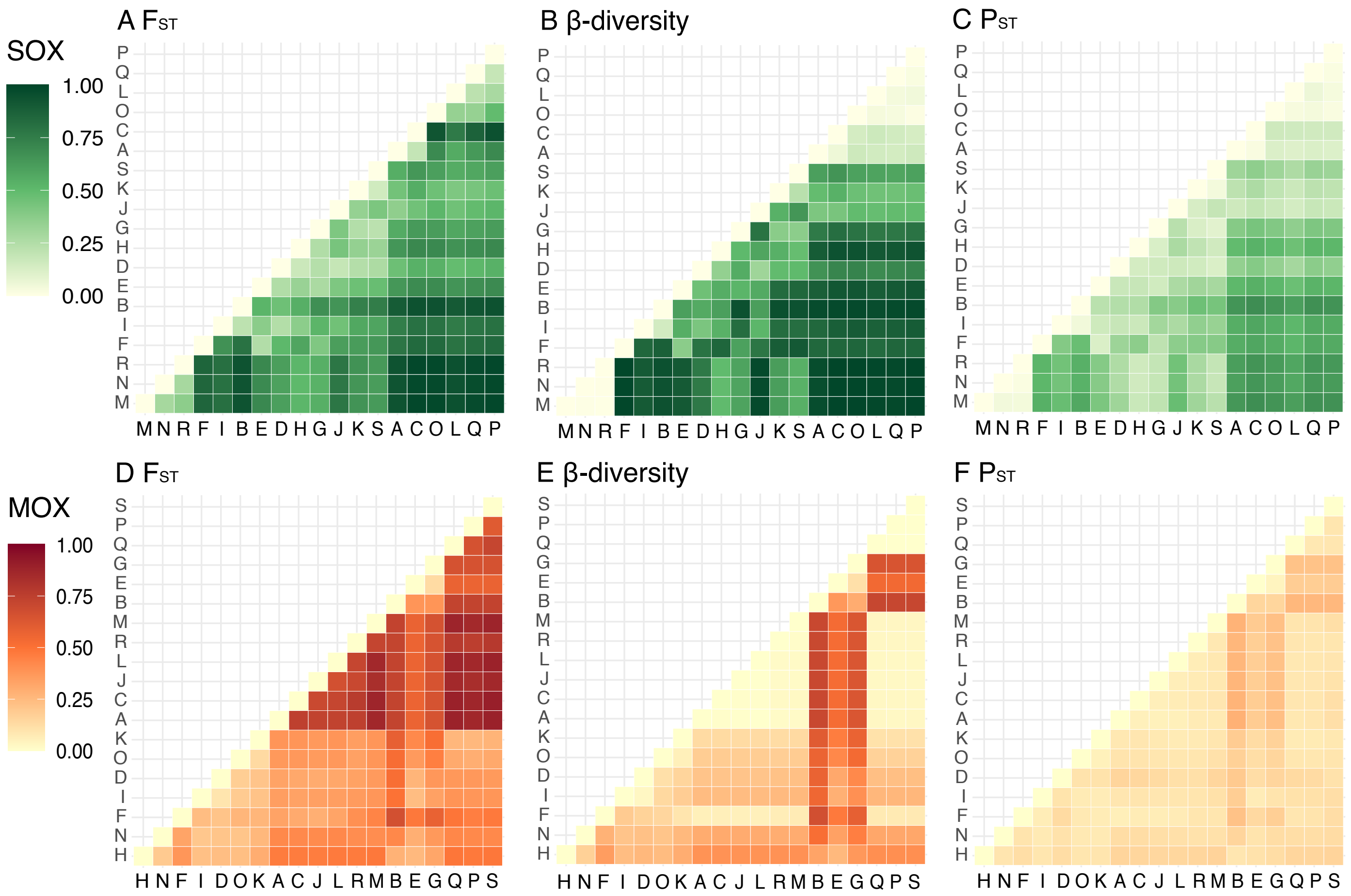


**Figure S5. Inter-sample diversity measured for A-C) SOX and D-F) MOX genomes.** Sample order is determined by hierarchical clustering based on $F_{ST}$.

**A**

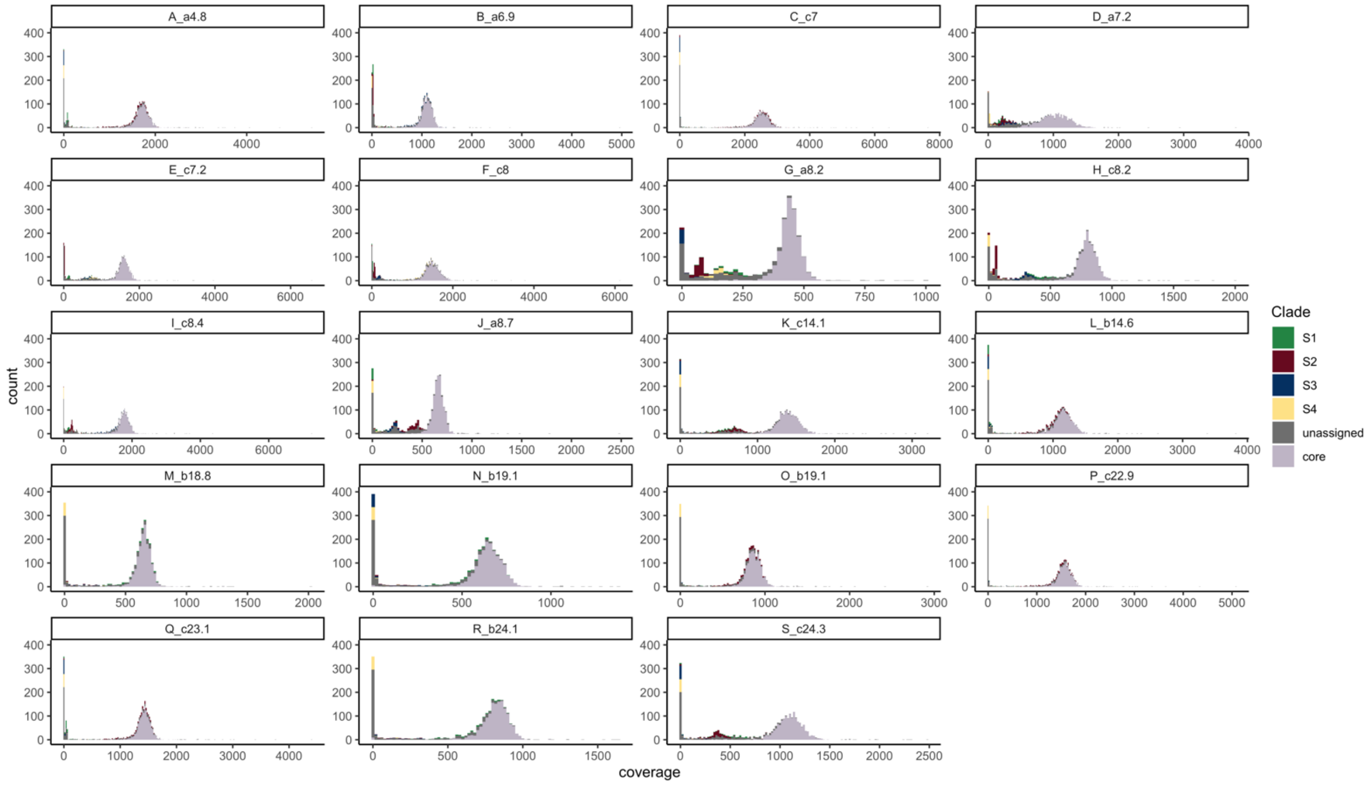


**B**

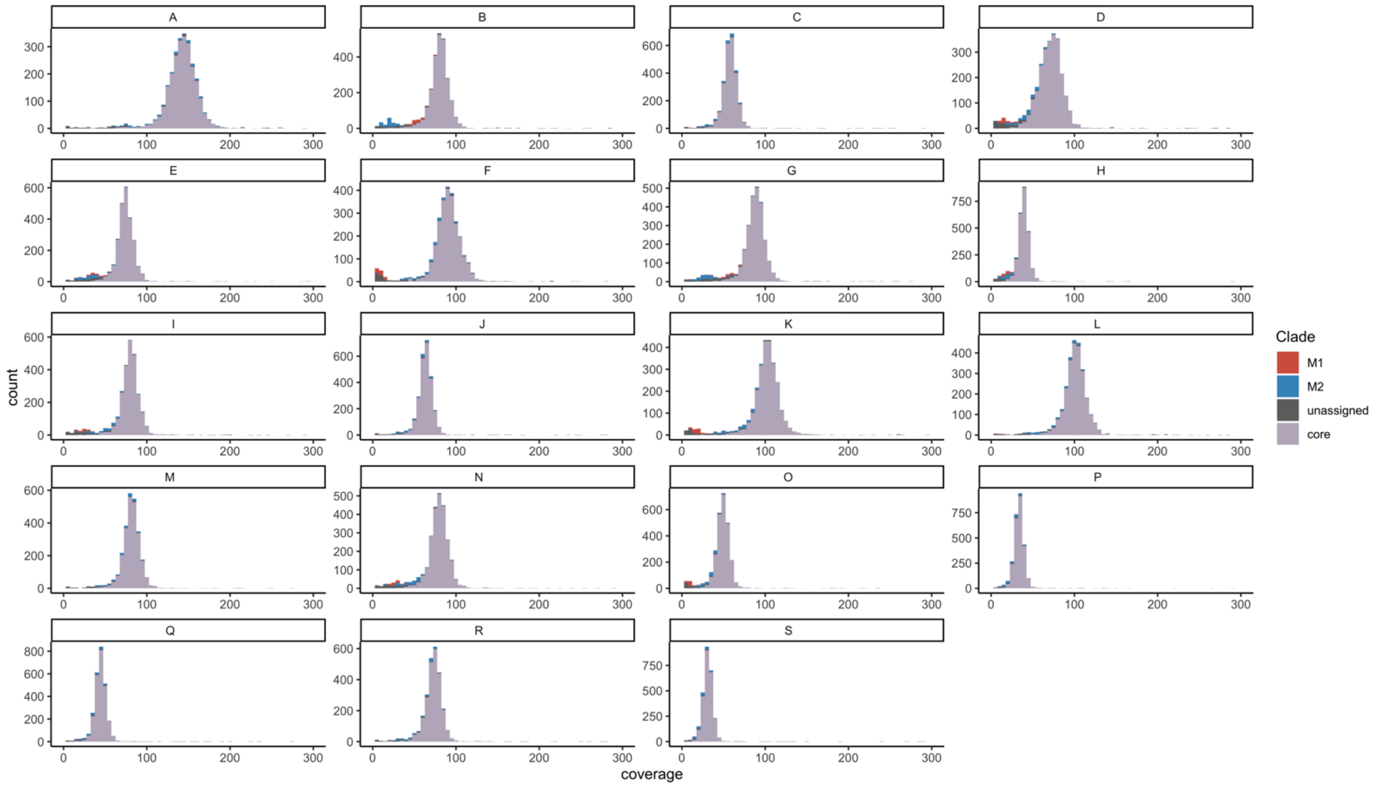


**Figure S6. Gene coverage distribution.** Stacked barplots showing gene coverage distributions for each individual sample for **A)** SOX and **B)** MOX pangenomes. Colors indicate core genes, strain-assigned accessory genes and unassigned accessory genes.

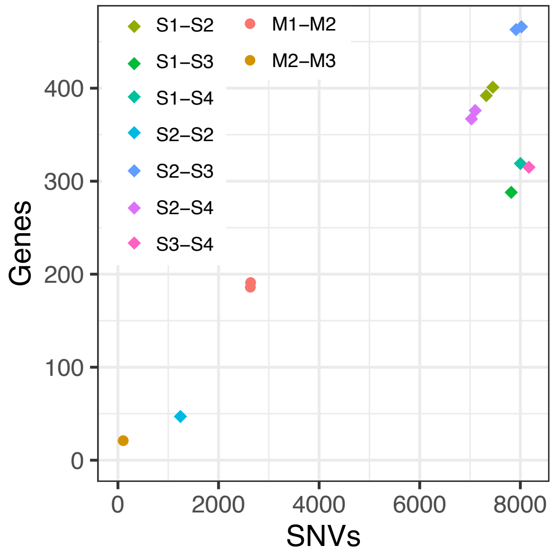


**Figure S7. Differentially present genes versus SNVs for reconstructed strains.** There is a high correlation between pairwise SNVs and differentially present genes, especially for MOX (Pearson's correlation coefficient, SOX $r^2$=0.717, MOX $r^2$=0.999). For SOX, intra-clade comparisons have ~8,000 SNVs but pairs that involve S2 have a higher number of differently present genes than pairs that do not involve S2. This difference can be traced back to the observation that more genes can be found in S2 but not in the other clade than vice versa (Table S4).

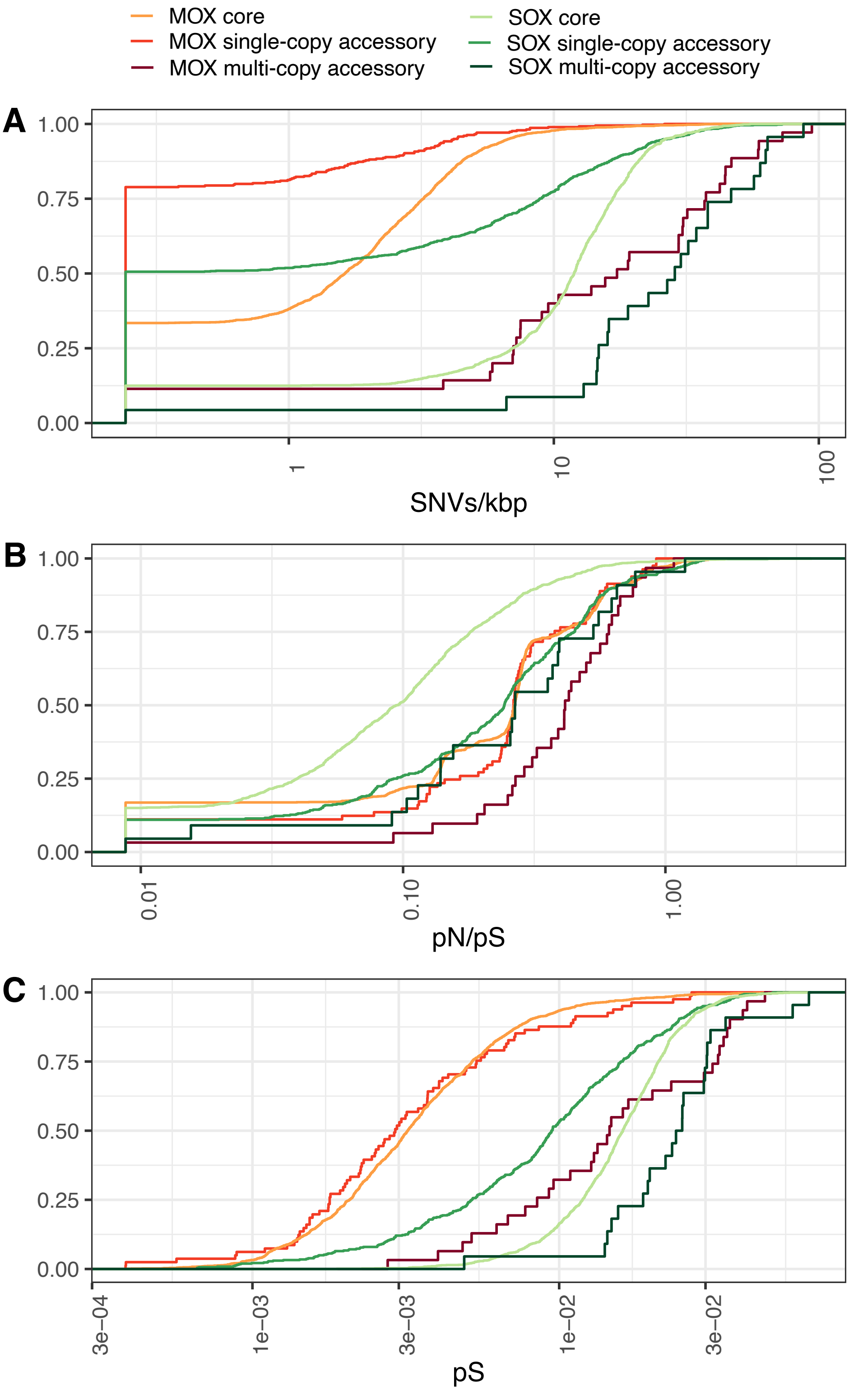


**Figure S8. Distribution of SNVs for core and accessory genomes in SOX and MOX. A**) Empirical cumulative distribution of the number of identified SNVs per kbp for SOX and MOX pangenomes. The distribution of SNVs/kbp is significantly different when comparing the SOX core and SOX accessory genomes and also when comparing the MOX core and MOX accessory genomes (Wilcoxon rank-sum test, p-value < $2.2 \times 10^{-16}$). Note that data points with SNVs/kbp = 0 are set to the minimum SNVs/kbp value found for representational purposes. **B, C)** Distributions of **B)** pN/pS and **C)** pS for SOX and MOX pangenomes. The pN/pS distribution is significantly different between SOX accessory genes and SOX core genes (Wilcoxon rank sum test p-value < $2.2 \times 10^{-16}$), where the median pN/pS is higher in the accessory genome. The pS distribution is significantly different between SOX accessory genes and SOX core genes (Wilcoxon rank sum test p-value < $2.2 \times 10^{-16}$), where the median pS is higher in the core genome. Neither the pN/pS nor the pS distributions of MOX core and accessory genes are significantly different (Wilcoxon rank sum test, p-values > 0.1). Note that multi-copy genes are excluded in this analysis and only genes with at least one SNV are included. Additionally, data-points with a pN/pS of 0 are set to the minimum pN/pS found.

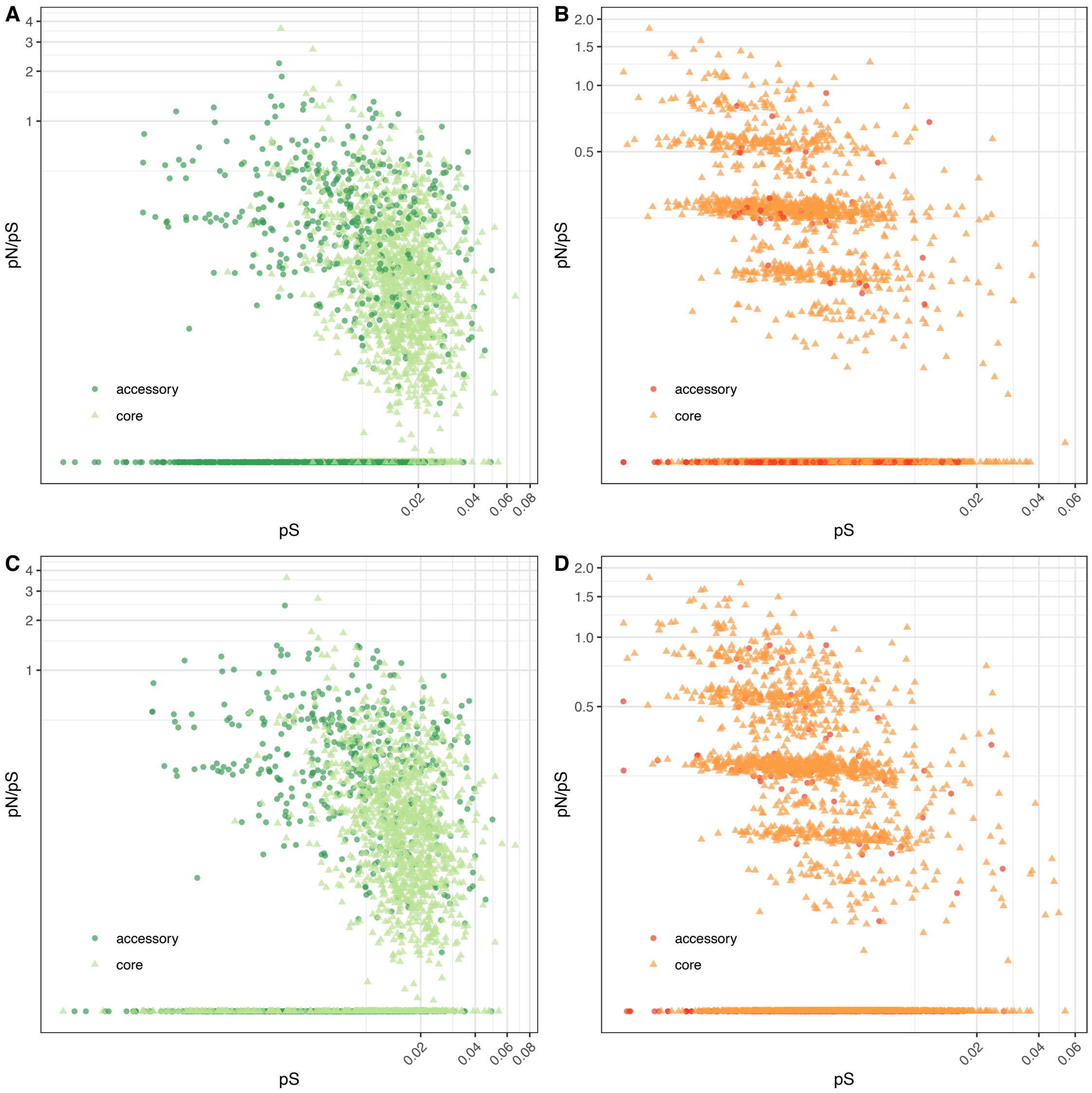


**Figure S9. pS and pN/pS distribution across core and single-copy accessory genes.** Distribution shown for **A)** SOX and **B)** MOX genomes considering SNVs across all 19 samples and for **C)** SOX and **D)** MOX genomes by considering SNVs called only in samples with dominant strains. Note that data-points with a pN/pS of 0 are set to the minimum pN/pS found for representational purposes.

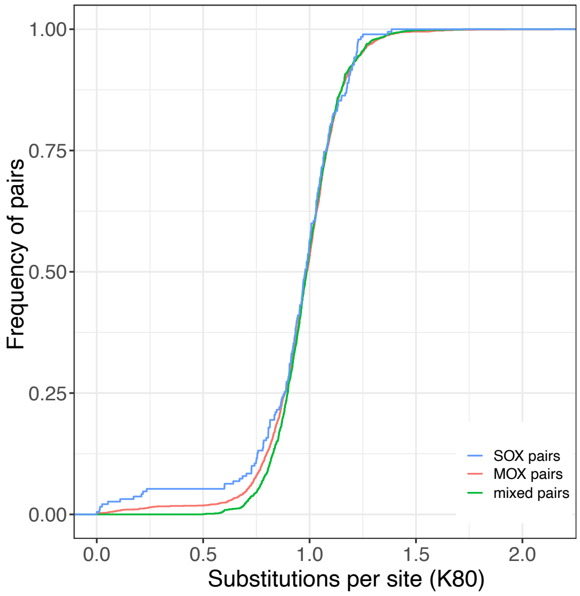


**Figure S10. Empirical cumulative distribution of transposase genetic diversity based on all pairs between 70 transposases in MOX and 20 transposases in SOX.**

### Supplementary tables

**Table S1. Pangenome reconstruction statistics across layers.**

| Layer | Number of genes | Median cluster size | Median sequence identity |
|---|---|---|---|
| **SOX** | | | |
| 0 | 1408 | 19 | 0.99 |
| 1 | 907 | 13 | 1 |
| 2 | 202 | 3 | 1 |
| 3 | 42 | 4.5 | 1 |
| 4 | 6 | 3 | 1 |
| 5 | 5 | 1 | 1 |
| Total | 2570 | | |
| **MOX** | | | |
| 0 | 2443 | 19 | 1 |
| 1 | 427 | 9 | 1 |
| 2 | 26 | 5 | 1 |
| 3 | 5 | 4 | 1 |
| 4 | 6 | 3.5 | 1 |
| Total | 2907 | | |

**Table S2. Statistics of MAGs (table in separate xlsx).** Completeness and contamination were estimated with CheckM.

**Table S3. Gene content intersections for the reconstructed strains.**

**A) SOX**

| **Strain** | **S1.4** | **S2.1** | **S2.2** | **S3.2** | **S4** |
|---|---|---|---|---|---|
| Samples | 3 | 2 | 4 | 2 | 1 |
| Total size | 293 | 391 | 398 | 295 | 322 |
| Overlap | 275 | 385 | 376 | 285 | |
| | 93.9% | 98.5% | 94.5% | 96.6% | |

| Gene content intersection | | | | | Size |
|---|---|---|---|---|---|
| | ■ | ■ | | | 144 |
| | | | ■ | | 71 |
| ■ | ■ | ■ | ■ | ■ | 67 |
| ■ | | | | | 65 |
| | ■ | ■ | | ■ | 61 |
| | | | | ■ | 59 |
| | | | ■ | ■ | 42 |
| ■ | ■ | ■ | | ■ | 34 |
| ■ | | | ■ | ■ | 33 |
| ■ | | | ■ | | 32 |
| ■ | ■ | ■ | | | 24 |
| | ■ | ■ | ■ | | 21 |
| | | ■ | | | 17 |
| ■ | ■ | ■ | ■ | | 14 |
| ■ | | | | ■ | 11 |
| | ■ | | | | 8 |
| | ■ | ■ | ■ | ■ | 6 |
| ■ | ■ | | | | 5 |
| | ■ | | | ■ | 4 |
| ■ | | ■ | | | 2 |
| | | ■ | ■ | | 2 |
| ■ | | ■ | ■ | | 2 |
| | | ■ | ■ | ■ | 2 |
| | ■ | | ■ | | 1 |
| ■ | ■ | | ■ | | 1 |
| ■ | ■ | | | ■ | 1 |
| ■ | | ■ | | ■ | 1 |
| ■ | | ■ | ■ | ■ | 1 |
| | | | | | 731 |

**B) MOX**

| **Strain** | **M1.2** | **M2.2** | **M2.3** |
|---|---|---|---|
| Samples | 1 | 5 | 4 |
| Total size | 144 | 198 | 203 |
| Overlap | | 157 | 151 |
| | | 79.3% | 74.4% |

| Gene content intersection | | | Size |
|---|---|---|---|
| | ■ | ■ | 113 |
| ■ | ■ | ■ | 77 |
| ■ | | | 65 |
| | | ■ | 12 |
| | ■ | | 7 |
| ■ | ■ | | 1 |
| ■ | | ■ | 1 |
| | | | 276 |

**A)** A total of 731 accessory genes could be assigned to five SOX strains across 12 samples. The gene content of clade S1 was reconstructed from strain S1.4 (dominant in samples M, N, R), the gene content of clade S2 was reconstructed from strain S2.1 (dominant in samples A, C) and strain S2.2 (dominant in samples L, O, P, Q), the gene content of clade S3 was reconstructed from strain S3.2 (dominant in samples B, I), and S4 was dominant in sample F. There are two strains for clade S2, total size: 418, intersection: 371 (88.8%). **B)** A total of 276 accessory genes could be assigned to 3 MOX strains across 10 samples. The gene content of clade M1 was reconstructed from strain M1.2 (dominant in sample

B) and the gene content of clade M2 was reconstructed from strain M2.2 (dominant in sample A, C, F, L, M) and M2.3 (dominant in sample K, P, Q, S). There are two strains for clade M2: total size 211, intersection: 190 (90.0%). Note that the gene content for each sample with a dominant strain has been reconstructed independently. Thus, the overlap of the reconstructed gene content of samples with the same dominant strain serves as a statistic to assess the robustness of the approach. For SOX, we find that more than 93% of strain-assigned genes are reconstructed across all samples containing that particular strain. In addition, the two strains of clade S2 overlap in 89% of their accessory genes. For MOX, we find that more than 74% of the strain-assigned genes are reconstructed across all samples that contain the strain and the two strains of clade M2 overlap in 90% of their accessory genes.

**Table S4. Pairwise differences for reconstructed strains.**

| Strain 1 | Strain 2 | Strain 1 \ Strain 2 | Strain 2 \ Strain 1 | # differential SNVs |
|---|---|---|---|---|
| SOX | | | | |
| S1.4 | S2.1 | 147 | 245 | 7323 |
| S1.4 | S2.2 | 148 | 253 | 7454 |
| S1.4 | S3.2 | 143 | 145 | 7819 |
| S1.4 | S4 | 145 | 174 | 8002 |
| S2.1 | S2.2 | 20 | 27 | 1243 |
| S2.1 | S3.2 | 281 | 185 | 8022 |
| S2.1 | S4 | 218 | 149 | 7029 |
| S2.2 | S3.2 | 283 | 180 | 7915 |
| S2.2 | S4 | 226 | 150 | 7103 |
| S3.2 | S4 | 144 | 171 | 8171 |
| MOX | | | | |
| M1.2 | M2.2 | 66 | 120 | 2633 |
| M1.2 | M2.3 | 66 | 125 | 2638 |
| M2.2 | M2.3 | 8 | 13 | 105 |

**Table S5. Putative phage contigs (table in separate xlsx).** "Coverage" gives the coverage as calculated with metaSPAdes. Contigs are assigned to a species, if their genes overlap with the species pangenome (for SOX and MOX) or with the species core genome (for MGS3). Only one contig has been detected as circular by VirSorter. Clustering showed 15 singletons and 12 clusters, where only the circular contig is grouping with additional phages from the database (cluster VC_68_0), thus, we identify it as a high-confidence phage with seven open reading frames. Based on vConTACT2, this phage clusters with *Alces alces* faeces associated microvirus MP12 5423, *Bdellovibrio* phage phiMH2K, *Chlamydia* phage 2, *Chlamydia* phage 4, *Chlamydia* virus CPAR39, Gokushovirinae Bog1183_53, Gokushovirinae Bog5712_52, Gokushovirinae Bog8989_22, Gokushovirinae Fen672_31, Gokushovirinae Fen7875_21, Guinea pig *Chlamydia* phage, and Marine gokushovirus. AAI based on one to three homologs is estimated between 35% (*Alces alces* faeces associated microvirus MP12 5423) and 47% (Gokushovirinae Bog1183_53). It has thus been termed *Gokushovirinae* sp. isolate VC_68_0 (GenBank accession OL437471.1) These phages are annotated as ssDNA viruses belonging to the subfamily Gokushovirinae. They have been mostly reconstructed from metagenomes or they have been isolated from Chlamydia. Here, we do not detect any additional MAG in sample E, which suggests that no abundant community member is present in this sample. Thus, the reconstructed phage could infect SOX or MOX or it could be a transient member in this community.

**Table S6. Functional annotation of A) SOX and B) MOX accessory genomes (table in separate xlsx).** The binary code represents whether a gene has been assigned to a specific strain. Additionally, the colors indicate strain- and clade-specific genes. Stars show genes that are identified as multi-copy, transposase, restriction-modification (RM) related or CRISPR-Cas related genes. One-letter abbreviations for the functional categories: A, RNA processing and modification; B, chromatin structure and dynamics; C, energy production and conversion; D, cell cycle control, cell division, chromosome partitioning; E, amino acid transport and metabolism; F, nucleotide transport and metabolism; G, carbohydrate transport and metabolism; H, coenzyme transport and metabolism; I, lipid transport and metabolism; J, translation, ribosomal structure and biogenesis; K, transcription; L, Replication, recombination and repair; M, cell wall/membrane/envelope biogenesis; N, cell motility; O, post-translational modification, protein turnover, and chaperones; P, inorganic ion transport and metabolism; Q, secondary metabolites biosynthesis, transport, and catabolism; S, no functional prediction; T, signal transduction mechanisms; U, intracellular trafficking, secretion, and vesicular transport; V, defense mechanisms; W, extracellular structures; X, mobilome: prophages, transposons; Z, cytoskeleton.

**Table S7. MOX and SOX orthologous gene pairs and their functional annotation (table in separate xlsx).**

**Table S8. Pfam accession numbers used in this analysis to infer genes related with Restriction-Modification systems and CRISPR-Cas (table in separate xlsx).**
